# A system biology-oriented investigation of Arabidopsis proteomes altered in chloroplast biogenesis and retrograde signaling reveals adaptive responses at whole cell level

**DOI:** 10.1101/2024.06.24.600381

**Authors:** Dario Di Silvestre, Nicolaj Jeran, Guido Domingo, Candida Vannini, Milena Marsoni, Stefania Fortunato, Maria Concetta de Pinto, Alberto Tamborrino, Yuri Luca Negroni, Michela Zottini, Lien Tran Hong, Andrea Lomagno, Pierluigi Mauri, Paolo Pesaresi, Luca Tadini

**Author notes:** these authors contributed equally to the manuscript (Dario Di Silvestre, Nicolaj Jeran, Luca Tadini).

## Abstract

Communication across different plant cell compartments relies on an intricate network of molecular interactions, required for the orchestration of organelle development and adaptation to the environment. In this scenario, the Pentatricopeptide Repeat (PPR) Protein GENOMES UNCOUPLED1 (GUN1) plays a key role in transferring information from both developing and mature chloroplasts to the nucleus with the aim to coordinate gene expression between the two genomes. However, its role and the related signaling molecules are still under debate. To help shed light on this matter, we attempted the holistic description of *Arabidopsis thaliana* proteome upon perturbation of chloroplast biogenesis by lincomycin (Lin), in a genetic context devoid of GUN1-dependent plastid-to-nucleus signaling pathway. Furthermore, the topological analysis of protein-protein interaction (PPI) and protein co-expression networks allowed the identification of protein hubs/bottlenecks characterizing genotypes and conditions, such as proteases, HSPs/Chaperones and redox proteins. Taken together, our findings indicate that GUN1 is required to orchestrate a plastid-located response to plastid protein synthesis inhibition while, in its absence, the reorganization of the activities associated with extra-plastid compartments, such as cytosol, vacuole and mitochondria, prevails. From this landscape, we documented a new role of the Oxygen Evolving Complex subunit PsbO, which appears to be an unconventional photosynthetic protein, as it accumulates in non-photosynthetic plastids and plays a central role in promoting chloroplast breakdown when plastid functions are altered.

## 1. Introduction

Understanding the mechanisms that underlie the communication among plant cell organelles and compartments will pave the way for elucidating the molecular mechanisms at the basis of plant adaptation to environmental changes, and for developing strategies to increase yield and stress tolerance (Wu & Bock, 2021). Signals between nucleus and organelles, i.e. anterograde and retrograde communication, are integrated into a complex network of interactions that exerts control over developmental stages, adaptation and/or the functional status of the organelles (Ng *et al*, 2014; Wu & Bock, 2021). In this scenario, chloroplasts and mitochondria, besides being central metabolic factories, are important players in sensing developmental and environmental stimuli and in conveying to the nucleus essential information for the biogenesis and maintenance of organelle functions (Wang *et al*, 2020). During the last two decades, the interest towards organelle-dependent anterograde and retrograde communication has increased, and substantial progress has been made in identifying key components and pathways. We learnt, for instance, that upon accumulation of unfolded or misfolded proteins in the organelles, mitochondrial (mtUPR) and chloroplast (cpUPR) <u>U</u>nfolded <u>P</u>rotein <u>R</u>esponse(s) are activated. In this frame, stressed mitochondria (Wang & Auwerx, 2017) and chloroplasts (Llamas *et al*, 2017) inform the nucleus of the need to increase the transcription of protective genes, including those encoding chaperones and proteases (Richter *et al*, 2023). In addition, several putative signaling molecules have been identified, such as carotenoids (Woodson *et al*, 2011; Moreno *et al*, 2021), metabolites (Milanesi *et al*, 2020; Fang *et al*, 2019; Vogel *et al*, 2014), intermediates of tetrapyrrole biosynthesis (Woodson *et al*, 2011), isoprenoid precursors (Xiao *et al*, 2012) nucleotides (Ono *et al*, 2021b; Estavillo *et al*, 2011) and redox signals (Jan *et al*, 2022; Maruta *et al*, 2012). From this basket of molecules, the GENOME UNCOUPLED 1 (GUN1) protein has been shown to be a key mediator of chloroplast retrograde communication through a model in which protein import capacity, folding stress and the cytosolic HSP90 complex control retrograde communication (Wu *et al*, 2019a; Tadini *et al*, 2020c). Other studies depicted GUN1 as a safeguard during the critical step of seedling emergence from darkness, by regulating transcription factors involved in light response, photomorphogenesis, and chloroplast development (Hernández-Verdeja *et al*, 2022; Veciana *et al*, 2022; Wu *et al*, 2019a), in the redox changes occurring during biogenic retrograde signaling (Fortunato *et al*, 2022), as well as in the acquisition of basal thermotolerance in *Arabidopsis thaliana* (Lasorella *et al*, 2022).

Although the direct communication between chloroplasts and mitochondria is still poorly understood, the evidence of common mitochondria- and plastid-to-nucleus retrograde pathways is widely supported (Wang *et al*, 2020). For instance, the existence of numerous chloroplast and mitochondrial dual-located proteins involved in diverse functions (Xu *et al*, 2013; Mitschke *et al*, 2009), together with the notion that chloroplasts and mitochondria have a synergistic effect in regulating nuclear gene expression (Pesaresi *et al*, 2006), strongly indicate the existence of a tightly integrated network of mitochondrial and chloroplast anterograde and retrograde signaling pathways (Wang *et al*, 2020). Noteworthy, organelle-to-nucleus retrograde communication also requires cytosolic- and nuclear-located components, such as kinases and transcription factors, which are shared between chloroplasts and mitochondria (Wurzinger *et al*, 2018; Blanco *et al*, 2014; Shapiguzov *et al*, 2019; De Clercq *et al*, 2013; Van Aken & Pogson, 2017; Van Aken *et al*, 2013), and whose stability is thought to be influenced by cytosolic folding stress (Wu *et al*, 2019a; Tadini *et al*, 2020a). These findings highlight the active roles of cytosolic and vacuolar compartments in the organellar quality check and degradation, orchestrated by retrograde signaling. Importantly, organelles dismantling triggers programmed cell death which, on one hand, is essential for plant development and response to biotic and abiotic stresses and, on the other hand, it ensures the maintenance of a healthy population of organelles while promoting nutrient re-distribution to sink tissues (Woodson, 2022; Van Aken & Van Breusegem, 2015).

The complexity and multitude of signals underlying the different communication pathways highlight how challenging it is to gain a clear picture of the molecular mechanisms responsible for organelle-nucleus and organelle-organelle communication. To shed further light on these sophisticated communication pathways and untangle the complex network of molecular relationships, we applied omics data-derived systems biology approaches (Di Silvestre *et al*, 2018). In the landscape of studies focused on investigating organelle signaling only a minority was based on omics data. Namely, some of them combined transcriptomic and proteomic profiles (Wu *et al*, 2019b; Marino *et al*, 2019), others took into consideration metabolomics (Bjornson *et al*, 2017), while recent studies were exclusively based on high-throughput proteomic analysis (Wu *et al*, 2018, 2019a; Tadini *et al*, 2020c). To take advantage of the plethora of information contained in -omics profiles, here we adopted a computational approach based on graph theory and network analysis, which represents a novel element in the study of plant retrograde signaling and chloroplast homeostasis. To exploit the correlation between the protein network’s structure and the supported functions, protein profiling and quantification were modeled as PPI and co-expression networks (Jeong *et al*, 2001; Vella *et al*, 2017). In previous studies, the functional and topological evaluation of these models was proven fruitful for the identification of proteins, defined as hubs or bottlenecks, as key players in the regulation of mechanisms underlying the investigated conditions (Di Silvestre *et al*, 2021, 2022). By following this strategy, in the present study we investigated at the system level the proteomes obtained from seedlings of *Arabidopsis thaliana* wild-type (Col-0) and *gun1-101* and *gun1-102* mutant plants treated or not with lincomycin (Lin), a chloroplast-specific translation inhibitor widely used to inhibit chloroplast biogenesis and to trigger the plastid-to-nucleus retrograde communication (Koussevitzky *et al*, 2007; Chotewutmontri & Barkan, 2018). Hypotheses and key players emerged from our holistic view, were validated at the experimental level. Our findings indicate that, in the presence of Lin, cell responses rely on the plastid compartment in a GUN1-dependent manner, while in the *gun1*-mutant background, the activity of extra-plastid compartments prevails. Moreover, results obtained by complementary approaches depict the Oxygen Evolving Complex subunit PsbO as an atypical Photosynthesis-related protein that accumulates in non-photosynthetic plastids and appears to have a superhub role in modulating chloroplast dismantling upon impairment of plastid functions.

## 2. Results

### 2.1 Proteome profiles of Arabidopsis Col-0 and *gun1* seedlings grown on lincomycin

To shed light on adaptive responses triggered by inhibition of plastid protein synthesis and chloroplast biogenesis, we combined and analyzed three independent proteome datasets (named Set1, Set2 and Set3) obtained from *Arabidopsis thaliana* Col-0 and *gun1* seedlings, grown on MS medium in absence and presence of Lin (-/+Lin) (**Figure 1 A**). A large number of proteins was identified in set1 (6852; this work) and set3 (7095; G., Z., Wu et al., 2019), containing proteins from total seedling extracts, while a slightly lower number was identified in Set2, which is enriched in soluble proteins from seedlings (5051; Tadini et al., 2020) (**Figure 1 B**). Globally, 9024 distinct proteins were identified by combining 36 LC-MS/MS total runs (Col-0, n = 9; *gun1*, n = 9; Col-0+Lin, n = 9; *gun1*+Lin, n = 9). A comparable number, about 7000, was found in each condition (**Figure 1 C** and **D**), while 837 proteins were in common among all the 36 samples analyzed (**Table S1**). Of note, about 30% of all identified proteins were characterized by an average spectral count (SpC) ≥ 1. The correlation among the global protein profiles indicated a higher divergence between proteomes of seedlings grown in the absence/presence of Lin in the MS medium (Col-0 *vs* Col-0+Lin, r = 0.84; *gun1 vs gun1*+Lin, r = 0.80), rather than between Col-0 and *gun1* genotypes (r = 0.98), as well as between +Lin growth conditions (r = 0.98) (**Figure 1 E**). A similar trend emerged when the differential enrichment of Gene Ontology (GO) biological process (BP) terms, such as Pigment metabolism, Photosynthesis, Metabolism, Transport, Organelle Organization, RNA processing/Translation, REDOX homeostasis and Development/Morphogenesis, were considered (**Figure 2, Table S2**). Most BPs were largely enriched in Col-0 and *gun1* seedlings extracts grown in control conditions (-Lin), with the only exceptions represented by Development/Morphogenesis, RNA processing/Translation and Transport processes, which were found to be enriched in seedlings grown in presence of Lin. In particular, Lin-mediated inhibition of plastid translation appears to stimulate the nuclear and intra-Golgi vesicle-mediated transport (**Figure 2 D**). Furthermore, BPs related to chloroplast organization and metabolism resulted down-regulated in favor of processes related to mitochondrial organization and ATP synthesis coupled to electron transport, indicating a possible role of mitochondria in compensating for chloroplast defects (**Figure 2 C**). These results agree with the most enriched GO molecular function (MF) terms, which mainly fell into RNA binding, REDOX activity, Folding/Response to stress, Tetrapyrrole binding, Isomerase activity and Endopeptidase activity. Of note, all MFs were most enriched in Col-0 and *gun1* genotypes with the exception of the Heat shock protein binding function, which characterized much more the *gun1*+Lin proteome (**Figure S1**).

**Figure 1.**
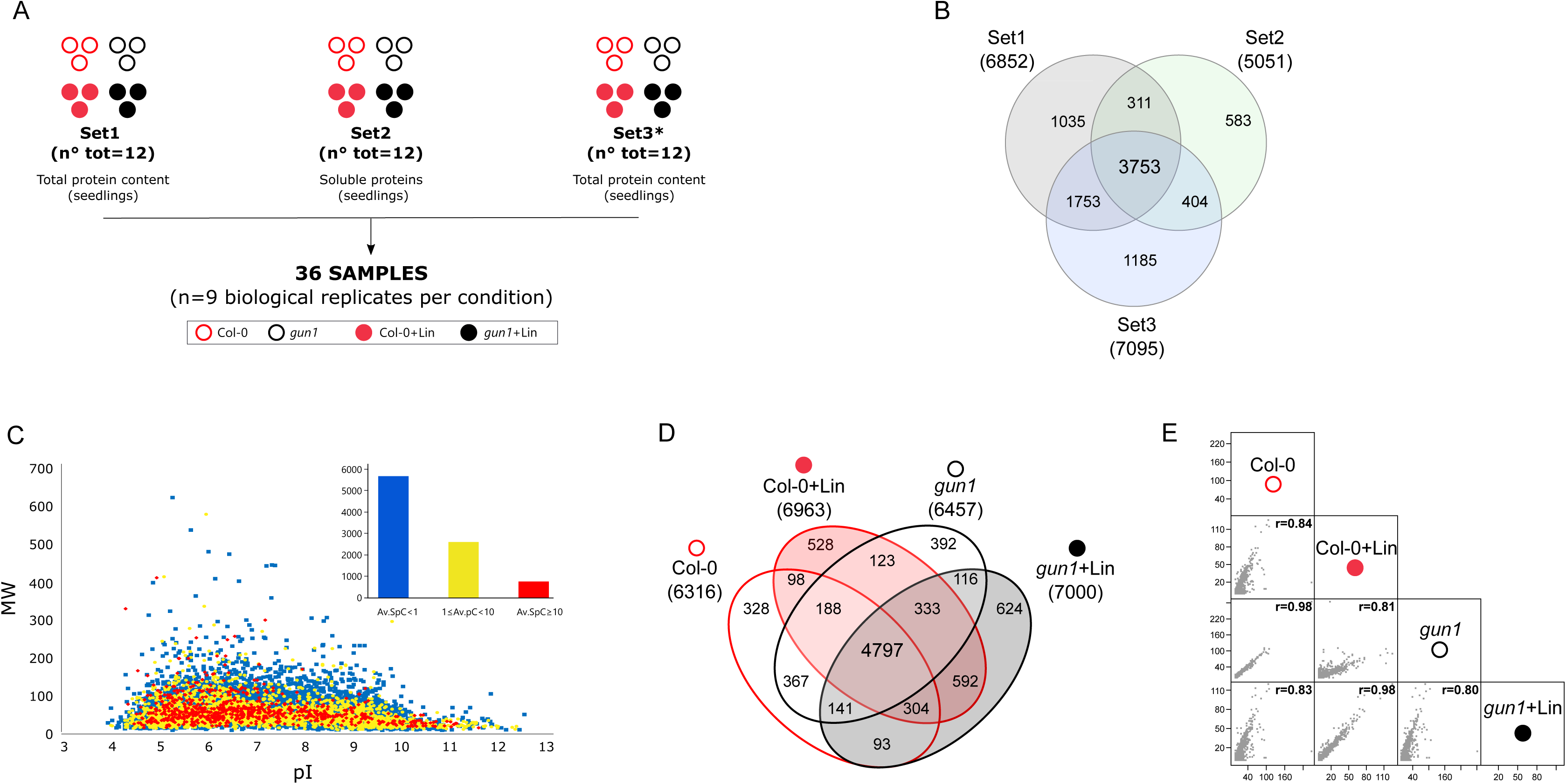
Proteomes from *Arabidopsis thaliana* Col-0 and *gun1* seedlings grown on MS medium in absence or presence (-/+) of Lin. **A**) A schematic overview of the experimental set-up exploited in this study, where conditions, datasets and replicates are shown. Each circle represents one replicate. Empty circles indicate control conditions whereas solid circles indicate Lin-grown samples. Col-0 is shown in red, *gun1* in black. **B**) Venn diagram of proteins identified in Set1, Set2 and Set3 (Set1: Col-0±Lin; *gun1-102*±Lin, 6 days old seedlings; our lab unpublished data; Set2: Col-0±Lin; *gun1-102*±Lin, 6 days old seedlings; (Tadini *et al*, 2020c); Set3, Col-0±Lin; *gun1-101*±Lin, 5 days old seedlings; (Wu *et al*, 2019a). **C**) 2D-Map (pI vs MW) of the total distinct identified proteins (n = 9024) and distribution of the average Spectral Count (Av.SpC) per protein (blue dots, Av.SpC < 1; yellow dots, 1 ≤ Av.SpC < 10; red dots, Av.SpC ≥ 10. **D**) Venn diagram of proteins identified in Col-0 and *gun1* mutant, grown or not with Lin (+/-Lin). **E)** Spearman’s correlation among Col-0, *gun1*, Col-0+Lin and *gun1*+Lin proteome profiles.

**Figure 2.**
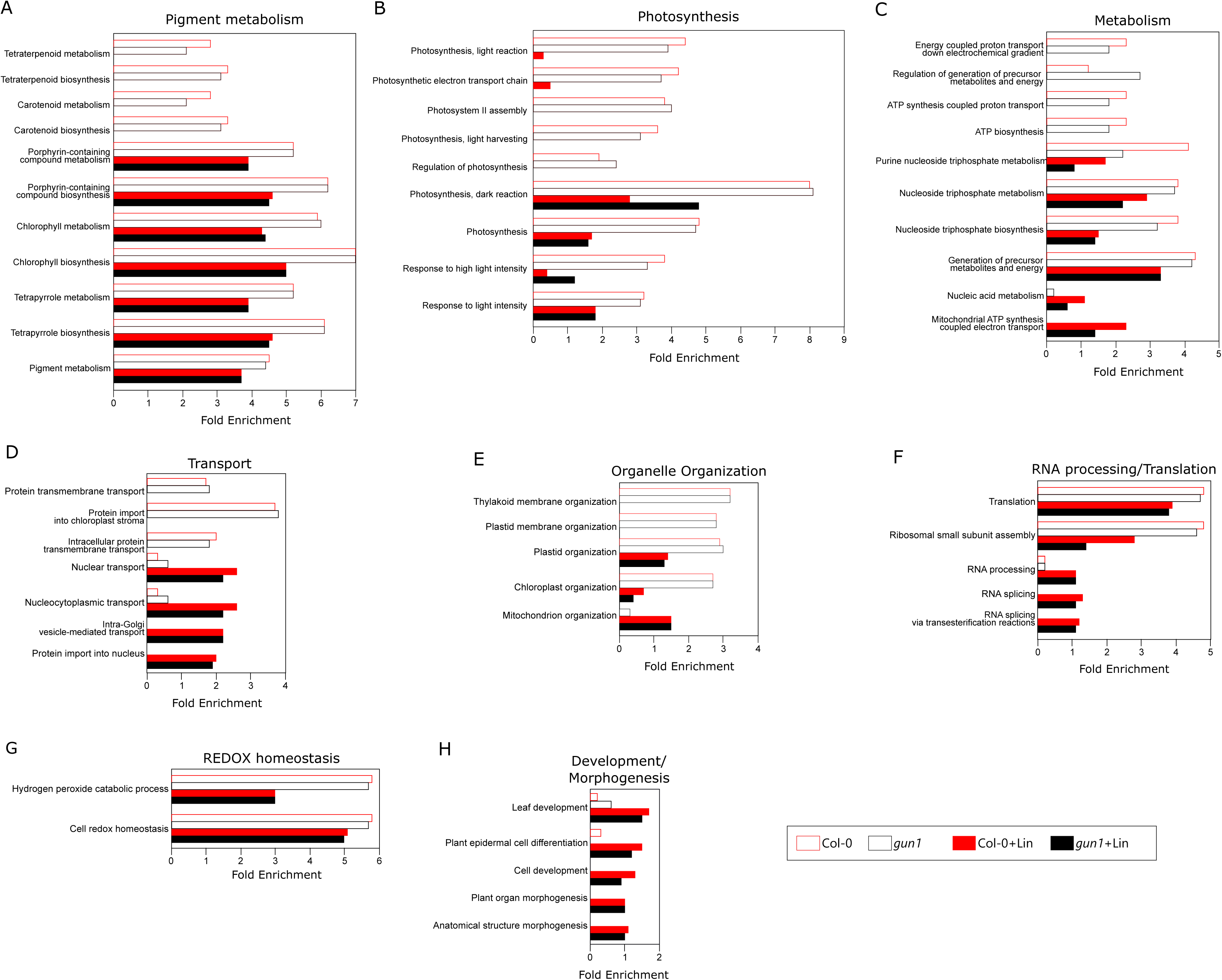
Biological processes (BPs) differentially enriched in *Arabidopsis thaliana* Col-0 and *gun1* seedlings, grown in absence or presence (-/+) of Lin. BPs were retrieved from TAIR and PANTHER databases, while those differentially enriched were extracted by using LDA (n = 9 per condition, p < 0.01, F ratio > 4.8). **A**) Pigment metabolism, **B**) Photosynthesis, **C**) Metabolism, **D**) Transport, **E**) Organelle Organization, **F**) RNA processing/Translation, **G**) REDOX homeostasis, **H**) Development/Morphogenesis.

### 2.2 Differentially Abundant Proteins in Col-0 and *gun1* seedlings grown on lincomycin

From the proteome profiles defined by Set1, Set2 and Set3 (**Figure 1**), 745 differentially abundant proteins (DAPs) were selected by a label-free approach based on Spectral Count (SpC) evaluation through Linear Discriminant Analysis (LDA, p≤0.01) (**Table S3)**. The largest differences were observed by comparing both Col-0 *vs* Col-0+Lin (537 DAPs) and *gun1 vs gun1*+Lin (547 DAPs), while far fewer DAPs were selected by comparing Col-0 *vs gun1* genotypes under similar growth conditions (-/+Lin), further supporting a larger effect due to the presence of Lin in the growth medium rather than the *gun1* mutation (**Figure 3 A**). A similar conclusion could be drawn from both Hierarchical Clustering and Principal Component Analysis (PCA), showing important differences between growth conditions (Col-0+Lin and *gun1*+Lin *vs* Col-0 and *gun1*) rather than genotypes (**Figure 3 B** and **C**). In addition, the good reproducibility of the protein expression patterns in the different biological replicate analyses is supported by the heat map where most DAPs were less abundant in plants grown in the presence of Lin. Furthermore, while PCA indicates that samples enriched in soluble proteins exhibit a slight deviation from the main group, the primary principal component (PC1), encompassing the highest variance at 56.1%, correlates with the distinct growth conditions (**Figure 3 C**).

**Figure 3.**
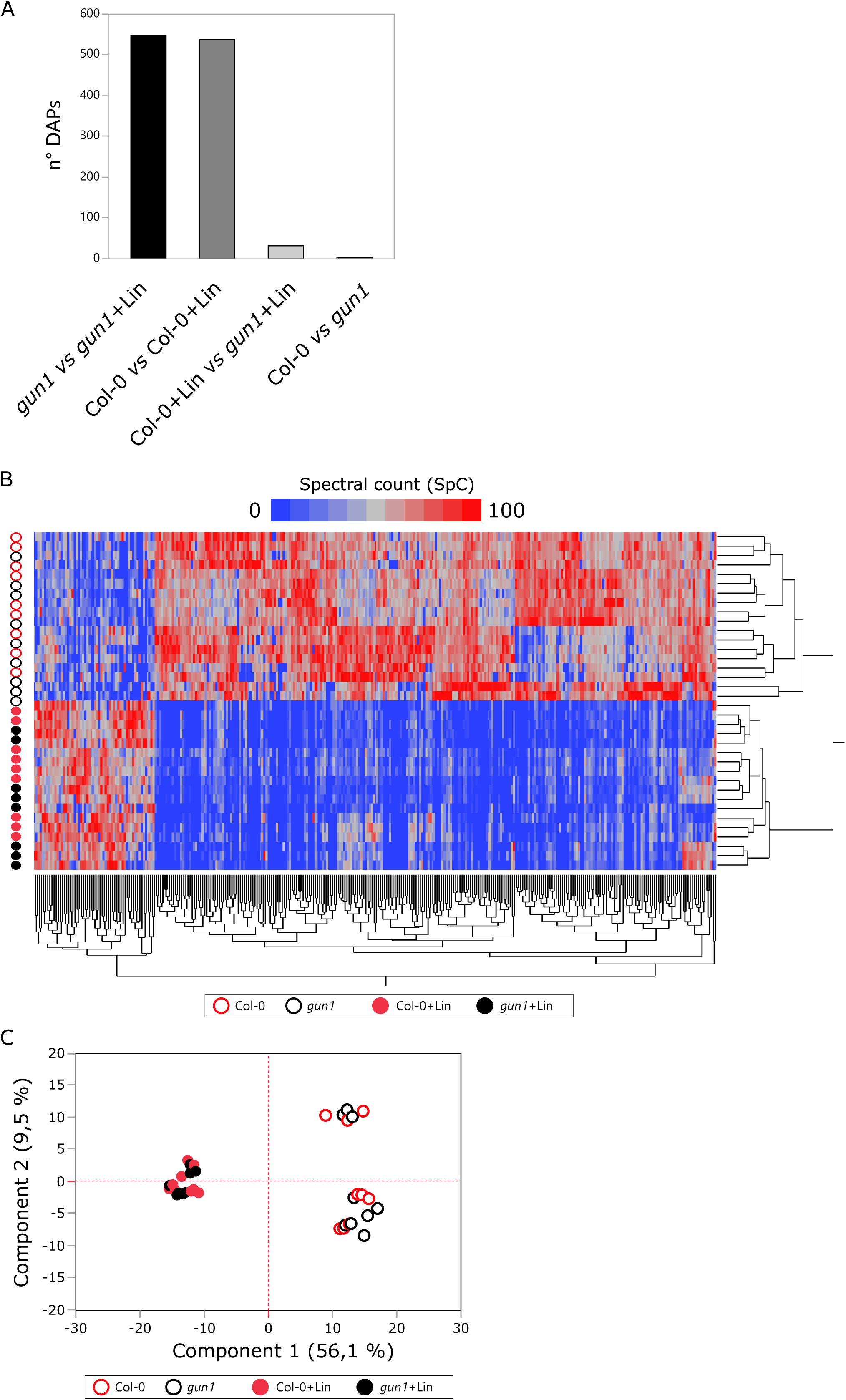
Differentially abundant proteins (DAPs) in *Arabidopsis thaliana* Col-0 and *gun1* seedlings, grown in absence or presence (-/+) of Lin in the MS medium. **A**) Histogram showing the number of DAPs (LDA, p ≤ 0.01) per comparison (*gun1 vs gun1*+Lin; Col-0 *vs* Col-0+Lin; Col-0+Lin *vs gun1*+Lin; Col-0 *vs gun1*). **B**) Hierarchical clustering and **C**) Principal Component Analysis (PCA) of DAPs selected with p ≤ 0.001.

To gain a comprehensive and in-depth representation of the proteins affected by the presence of Lin in the growth medium and/or by *gun1* mutation, high-confidence DAPs (p≤0.001, n=326) were modeled as PPI network and grouped in 41 functional modules representing pathways, GO biological processes (BPs), GO molecular functions (MFs), GO cellular components (CCs) and protein families (**Figure S2**). As already evidenced by the heat map (**Figure 3 B**), the protein abundance of most functional modules collapsed in both Col-0 and *gun1* genotypes upon plastid translation inhibition, i.e. in presence of Lin. They include proteins of photosystem II (PSII) and photosystem I (PSI) complexes, 30S and 50S ribosomal subunits, enzymes involved in tRNA aminoacylation, Calvin cycle, amino acid metabolism, ATP synthesis and chlorophyll biosynthesis, subunits of NAD(P)H dehydrogenase complex and proteins with peptidyl-prolyl cis-trans isomerase activity. Conversely, functional modules enriched in the TCA cycle, lipid metabolism, carbohydrate metabolism, protein folding, ubiquitination and proteolysis were most represented (**Figure S2**).

Since the reduced accumulation of large protein complexes and metabolic pathways are easily interpretable as the obvious consequence of Lin-mediated inhibition of protein synthesis and chloroplast biogenesis, we mainly devoted our attention to proteins whose abundance was retained or increased following growth on Lin-containing medium. Their classification in sub-categories, such as proteolysis, protein folding, oxidative stress response, fatty acid metabolism, ER-Golgi and organelle transport, allowed to highlight the enrichment of specific protein families, including Serine Carboxypeptidase (SCPL) in the Peptidases module (Huang *et al*, 2008; Havé *et al*, 2018) and the RPT/RPN subunits of 26S Proteasome in the Proteases module, especially in *gun1*+Lin (**Figure 4**). Moreover, some proteins were exclusively detected in the proteomes of Col-0 and *gun1* seedlings grown in the presence of Lin. These include the Acetyl-CoA Carboxylase 2 (ACC2) (Parker *et al*, 2014, 2016; Wang *et al*, 2018; Perez de Souza *et al*, 2020) involved in fatty acid metabolism, the Serine Carboxypeptidase-like 9 (SCPL9), involved in peptidase activity, and the nuclear pore complex protein NUP62 (Kemp *et al*, 2015; Zhao & Meier, 2011), involved in nuclear protein transport.

**Figure 4.**
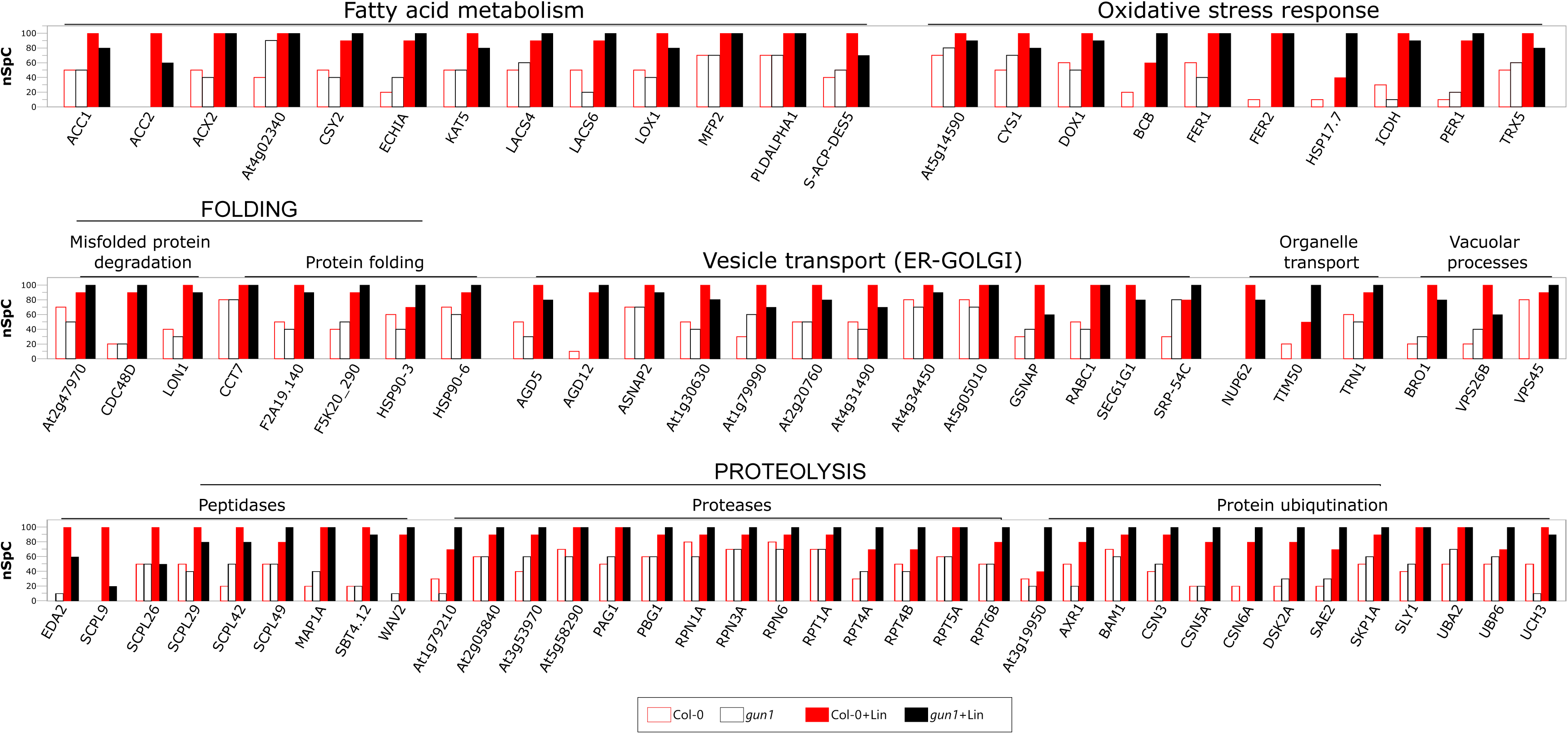
Proteins with increased abundance upon seedling growth in presence of Lin and functionally involved in Fatty acid metabolism, Oxidative stress response, Protein folding, Vesicle transport (ER-Golgi), Organelle transport, Vacuolar processes, and proteolysis. For each protein, the spectral count value normalized (nSpC) in the range 0-100 is shown (LDA, p ≤ 0.01).

### 2.3 Hubs and superhubs from PPI and co-expression network models

To gain new insights, in addition to DAPs, the characterized proteome profiles were modeled as PPI and protein co-expression networks, to be processed at the topological level and to identify hubs featuring the investigated genotypes/growth conditions. Following the combination between the characterized proteomes (Col-0, *gun1*, Col-0+Lin and *gun1*+Lin) and the interactome of *Arabidopsis thaliana*, a fully connected PPI network per condition was built. The four models showed scale-free distribution. Protein hubs were selected by combining betweenness, centroid and bridging centralities, and their significance was assessed by random networks that showed average betweenness values different from those calculated in the reference ones (**Figure S3 A**). Globally, 66 proteins were defined as hubs, 25 of them in all conditions, while 41 were genotype/growth condition-specific (**Figure 5 A**, **Table S4**). In the case of *gun1* seedlings, and especially in *gun1*+Lin seedlings, protein hubs are involved in the mitochondrial respiratory chain, ubiquitination, transcriptional regulation or belong to the copper amine oxidase family. On the other hand, photosynthesis and vacuolar-related proteins, including the vacuolar ATPase complex, were hubs mostly featuring Col-0+Lin seedlings. Noteworthy, the double relevance of proteins found to be both hubs and differentially abundant also emerged. Among others, Acetyl-CoA Carboxylase 1 (ACC1), involved in fatty acid biosynthesis (Damiano *et al*, 2018), was a hub in both genetic backgrounds under Lin-growth conditions and up-regulated in the same conditions. PsbO1, a subunit of the Oxygen Evolving Complex (Lundin *et al*, 2007; Jiang *et al*, 2017) behaved as a hub in both Col-0 and *gun1* seedlings and was down-regulated upon growth in Lin-containing medium. Moreover, Nitrate reductase [NADH] 2 (NIA2), involved in nitrate assimilation and detoxification of Reactive Oxygen Species (ROS) and Reactive Nitrogen Species (RNS) (Manbir *et al*, 2022; Khator & Shekhawat, 2020; Pan *et al*, 2019), was a hub in *gun1*+Lin seedlings. Intriguingly NIA2, down-regulated in Col-0+Lin, was unaltered in *gun1*+Lin when compared to the untreated controls (**Table S1**).

**Figure 5.**
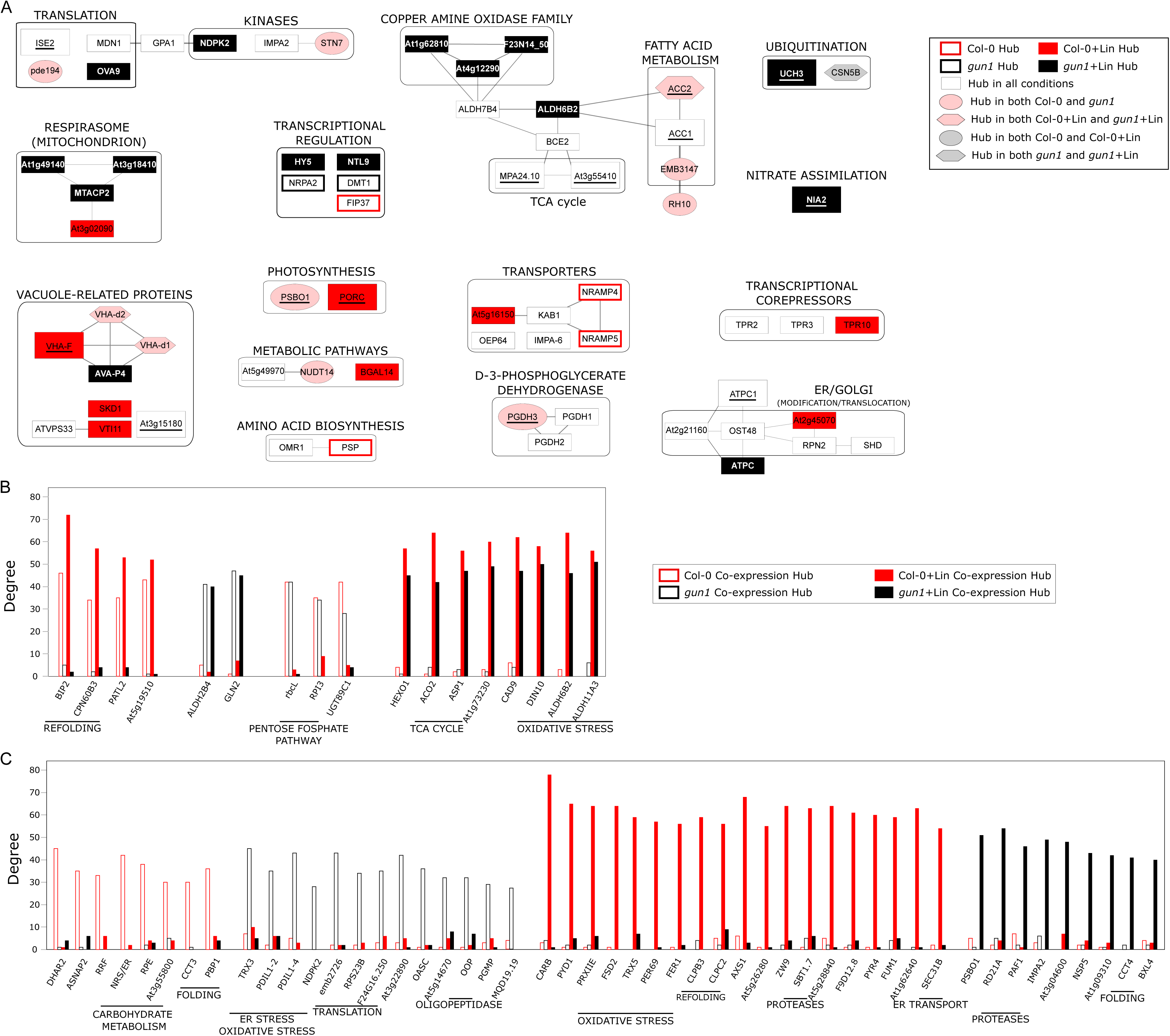
Hubs and superhubs from PPI and co-expression network models. **A**) PPI network hubs in *Arabidopsis thaliana* Col-0 and *gun1* seedlings, grown in absence or presence of Lin (-/+Lin); PPI hubs were selected taking into consideration betweenness, centroid and bridging centralities. Bigger and underlined nodes indicate proteins defined as hubs and differentially abundant (p ≤ 0.01), too. **B**) Differentially correlated proteins based on *gun1* mutation and Lin presence in the growth medium. **C**) Proteins highly correlated in Col-0, *gun1*, Col-0+Lin or *gun1*+Lin co-expression network models.

A further implementation of the hub set was achieved by integrating the topological analysis of the co-expression networks. Starting from the 837 proteins identified in all analyzed samples, we reconstructed a model per genotype/growth condition with the purpose of identifying proteins differentially correlated (**Table S5**). We assumed that the defined co-expression hubs could play a key role in regulating the mechanisms underlying the investigated conditions. Similarly to PPI hubs, their significance was assessed through the degree of centrality, calculated using random networks **(Figure S3 B)**. Upon growth on MS medium containing lincomycin, a high degree of correlation was observed in proteins involved in oxidative stress [dark inducible 10 (DIN10)], aldehyde dehydrogenase 6B2 (ALDH6B2), aldehyde dehydrogenase 11A3 (ALDH11A3), peroxiredoxin-II-E (PRXIIE), Fe superoxide dismutase 2 (FSD2), thioredoxin h-type 5 (TRX5), peroxidase69 (PER69), ferritin 1 (FER1) and TCA cycle (**Figure 5 B** and **C**). On the contrary, proteins involved in the Pentose Phosphate Pathway were highly correlated in Col-0 and *gun1* seedlings. Moreover, proteins involved in protein refolding, such as Heat shock 70 kDa proteins (BIP2; Srivastava et al., 2013; Yang et al., 2014), Chaperonin-60beta3 (CPN60B3; Klasek et al., 2020), Clp protease subunit ClpB homolog 3 (ClpB3; Llamas et al., 2017; Parcerisa et al., 2020), chaperone protein ClpC2 (Park & Rodermel, 2004) were highly correlated in Col-0 and especially in Col-0+Lin. Conversely, proteins involved in ER and oxidative stress [thioredoxin H-type (TRX3)], protein disulfide isomerase 1-2 (PDIL1-2) and protein disulfide isomerase 1-4 (PDIL1-4) were specifically high-correlated in *gun1* plants **(Figure 5 C)**.

The degree of correlation was also evaluated among proteins that physically interact and are involved in the same biological process, molecular function, or protein complex, confirming the observations emerged by quantitative analysis and PPI network models. In fact, in both Col-0+Lin and *gun1*+Lin proteomes, we highlighted an increasing correlation between proteins involved in protein folding, amino acid metabolism, fatty acid metabolism and peptidases. We also observed that the correlation of 14-3-3 proteins and ribosomal subunits was mainly affected by the lack of GUN1 protein (**Figure S4**). As expected, the correlation among Calvin-Benson cycle proteins collapsed in Col-0 and *gun1* seedlings +Lin, while proteins involved in ER stress were found specifically correlated in *gun1* genotype (**Figure S4**). On the basis of these observations, we selected a group of proteins, defined as superhubs, since they are topologically relevant in both PPI and co-expression models (**Table 1**). Among them, Oxygen-evolving enhancer protein 1-1 (PsbO1), involved in water oxidation at the Photosystem II, and 2-oxoglutarate dehydrogenase (At3g55410), involved in mitochondrial TCA cycle, were also found to accumulate differentially, supporting a role as key players in regulating the organelle adaptive responses.

**Table 1.**
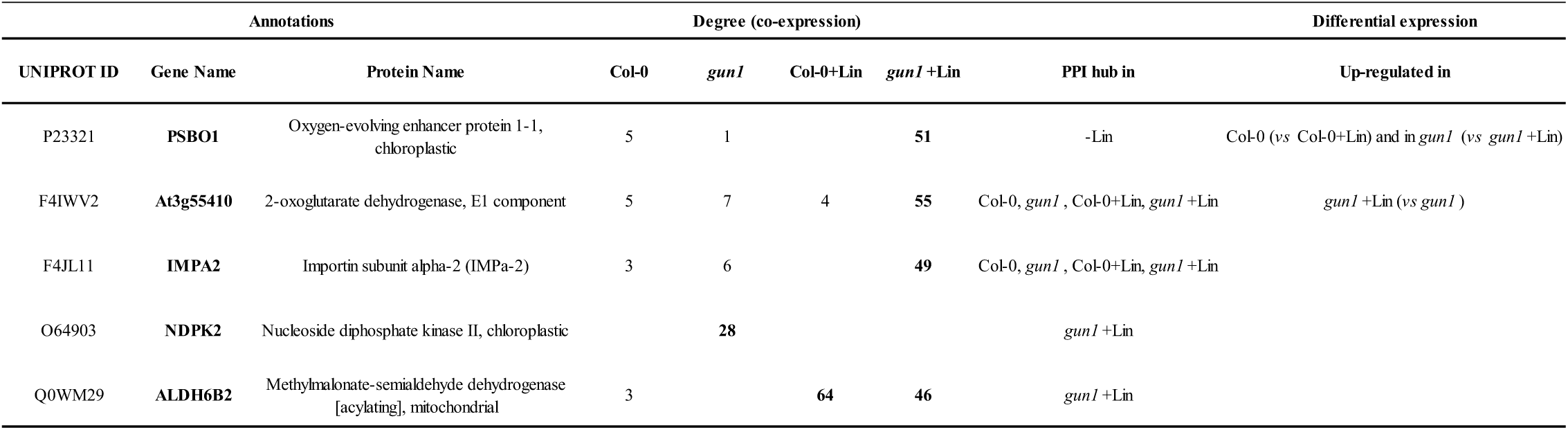
List of proteins elevated to hubs in both PPI and Co-expression network models and defined as superhubs.

### 2.4 Hub functional validation from PPI and co-expression networks: the extra-plastid compartment(s)

To functionally validate the computational analyses described above and to provide a wider picture of cellular functions with a potentially key role in the different conditions, a set of hub proteins highlighted by PPI and co-expression networks was taken into consideration for further investigations. First, the increased cytosolic folding stress, observed in *gun1*+Lin samples, was considered. Unlike most chloroplast-related proteins, those involved in ubiquitination processes were found to accumulate in Col-0 and *gun1* Lin-treated samples, together with proteins involved in plastid proteolysis. In particular, DAPs included peptidases, proteases and ubiquitination-related processes in +Lin samples (**Figure 4** and **Table S3**). Moreover, PPI networks highlighted a central topological role of Ubiquitin Carboxyl-terminal Hydrolase 3 (UCH3) protein in *gun1*+Lin samples (**Figure 5 A**). Similarly, the RESPONSIVE TO DEHYDRATION 21A (RD21A) protein, involved in cytosolic protein degradation and vacuolar-stress-mediated cell death (Koh *et al*, 2016; Boex-Fontvieille *et al*, 2015; Wang *et al*, 2008), was found as a co-expression hub only in *gun1*+Lin (**Figure 5 C**). Accordingly, UBQ11-specific immunoblot on total protein seedling extracts revealed a higher amount of ubiquitinated proteins in *gun1-102* samples grown in presence of Lin, with respect to Col-0+Lin, Col-0 and *gun1-102* samples (**Figure 6 A**). Similarly, the cytosolic chaperones HSP90s accumulated at higher levels in *gun1-102*+Lin seedlings (**Figure 6 A**), as described previously (Wu *et al*, 2019a; Tadini *et al*, 2020c). This observation indicated highly active cytosolic 26S proteasome-mediated protein degradation which was measured as the release of amino-methyl-coumarin from the fluorogenic substrate Suc-LLYY-NH-AMC (**Figure 6 B**). Compared to the other genotypes and conditions, proteasome activity was higher in *gun1-102*+Lin samples further corroborating our preliminary findings and indicating an increased cytosolic proteolysis (**Figure 4** and **Figure S4**).

**Figure 6.**
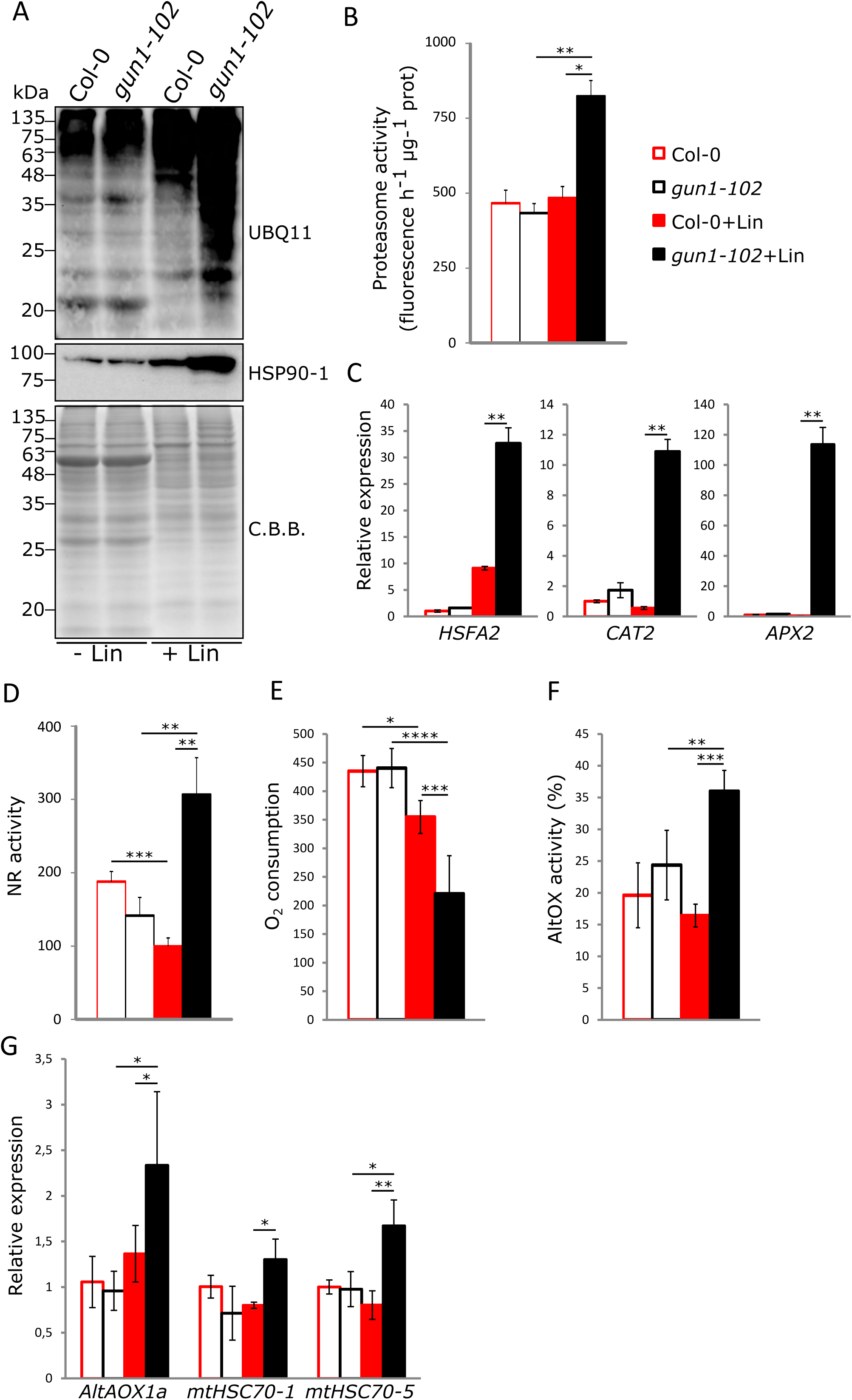
Validation of extra-plastid protein hubs. **A**) Cytosolic folding stress and ubiquitination of total protein extract in Col-0 and *gun1-102* samples grown in absence or presence of Lin (-/+Lin), shown by immunoblot analyses (UBQ11 and cytosolic AtHSP90-1 specific antibodies). Coomassie Brilliant Blue (C.B.B.) staining of SDS-PAGE is shown as loading control. **B**) Proteasome activity assay expressed as the fluorescence released by amino-methyl-coumarin. **C**) Real-time quantitative PCR on cytosolic ROS and stress-related marker genes *HSFA2*, *CAT2* and *APX2*. **D**) Nitrate reductase enzyme activity (NR). **E**) Measurement of Oxygen consumption (nmol O_2_/min/mg protein) by mitochondrial respiration in Col-0 and *gun1-102* -/+Lin samples, together with **F**) mitochondrial Alternative Oxidase (AltOX) activity. **G**) Real-time quantitative PCR on *AltOX1a*, *mtHSC70-1* and *mtHSC70-5* transcripts, used as molecular marker genes to investigate mitochondrion-to-nucleus retrograde signaling and mtUPR. Asterisks indicate significant differences (* p-value < 0.05; ** p-value < 0.01; *** p-value < 0.001; **** p-value < 0.0001).

To elucidate the contribution of Col-0 and *gun1* cytosolic compartments in the adaptation to the presence of Lin, the accumulation of *HSFA2* transcripts, a transcription factor with a key regulatory role in the activation of the cellular response upon several stresses (Nishizawa *et al*, 2006), and the expression of *CAT2* and *APX2* genes, encoding a catalase and an ascorbate peroxidase active in the peroxisome and cytosol, respectively, (Ono *et al*, 2021a; Fryer *et al*, 2003; Anjum *et al*, 2016) were analyzed by RT-qPCR performed on total RNA extracts from 6 days after sowing (DAS) seedlings (**Figure 6 C**). *HSFA2* expression was induced by Lin in Col-0 and at a higher level in *gun1-102* plants, consistent with the higher cytosolic folding stress observed in mutant plants (Tadini *et al*, 2020c; Wu *et al*, 2019a). More interestingly, *CAT2* and *APX2* gene expressions were largely up-regulated in *gun1-102* seedlings when grown with Lin, whereas no induction was detected in Col-0 seedlings grown in the same conditions, as reported previously (Fortunato *et al*, 2022).

Additionally, NIA2 involved in nitrogen assimilation by reducing nitrate, was a PPI hub in the *gun1*+Lin condition (**Figure 5 A**). Of note, the topological relevance emerged for NIA2 was in line with the nitrate reductase (NR) activity measured in our plants. As a matter of fact, upon growth on Lin, NR activity decreased in Col-0 seedlings, compared to untreated samples, while markedly increased in *gun1-102* (**Figure 6 D**). In addition to assimilating nitrate, NR can participate in the generation of Nitric Oxide (NO), known to induce the upregulation of the alternative oxidase (AltOX) upon stress conditions (Huang *et al*, 2002; Salgado *et al*, 2013). The mitochondrial compartment appeared to play a critical role in *gun1*+Lin samples, rather than in the other conditions, as indicated by the presence of “TCA cycle” and “Mitochondrion respiration” GO categories in the PPI network and co-expression analyses (**Figure 2**, **Figure 5**, **Figure S2**, **Figure S4**). NO serves as a crucial signaling molecule, yet its reversible interaction with COX in mitochondria can lead to detrimental effects by inhibiting respiratory chain activity (Wulff *et al*, 2009; Millar & Day, 1996).

Based on the insights depicting a possible interaction between NR activity and mitochondria, the O_2_ consumption and the AltOX activity were measured (**Figure 6 E, F**). While O_2_ consumption was similar between Col-0 and *gun1-102* seedlings under control conditions, respiration decreased in Col-0 and more markedly in *gun1-102* seedlings in the presence of Lin. In terms of AltOX activity, no differences were detected between untreated wild-type and *gun1-102* seedlings, and no changes were observed in Col-0 upon Lin treatment, while AltOX activity increased in *gun1-102*+Lin seedlings. Consistently, the expression of *AltOX1a* was increased in the presence of Lin only in *gun1-102* background (**Figure 6 G**), together with transcripts of nuclear genes, such as *mtHSC70-1* and *mtHSC70-5*, encoding mitochondria-located proteins involved at different levels in stress-response (**Figure 6 G**), indicating that the mitochondrial adaptive response also requires the mitochondria-to-nucleus retrograde communication. Noteworthy, the mitochondrial adaptive response appears to be triggered by the Lin-specific inhibition of plastid protein synthesis, as no major alterations of mTP-YFP mitochondria morphology were detected among the different genotypes by confocal microscopy fluorescent imaging (**Figure S5 A**). Overall, these observations indicate that upon impairment of plastid functions and in the absence of GUN1-mediated plastid-to-nucleus retrograde communication, mitochondria play a central role in cell detoxification together with vacuolar and cytosolic compartments.

### 2.5 Functional validation of hubs from PPI and co-expression networks: the plastid compartment

Many plastid-located proteins were also present among the identified PPI and co-expression hubs (**Table S4**, **Table S5**). In particular, 11 out of 19 co-expression hubs in Col-0+Lin seedlings were plastid-located proteins, indicating that upon Lin-inhibition of plastid translation, plant cell responses mainly rely on GUN1-mediated plastid activities. The main hubs that are activated following the inhibition of plastid protein translation include proteins involved in stress response mechanisms such as ROS scavenging and protein folding/degradation functions (PRXIIE, FSD2, ClpB3 and ClpC2). On the contrary, the superhub PsbO1 was the only plastid-located co-expression hub specific for *gun1*+Lin seedlings (**Figure 5 C and Table S5**). This observation is rather peculiar since PbsO1 is part of the Oxygen Evolving Complex (OEC) bound to Photosystem II and accumulates in plastid devoid of the photosynthetic apparatus (**Figure 7 A**, **Table S3**), unlike most photosynthesis-related proteins, including the other subunits of OEC (PsbQ, PsbR; **Figure 7 A**, **Table S3**) that have been reported to be either degraded in the cytosol as precursors or translationally inhibited (Wu *et al*, 2019a; Tadini *et al*, 2020c).

**Figure 7.**
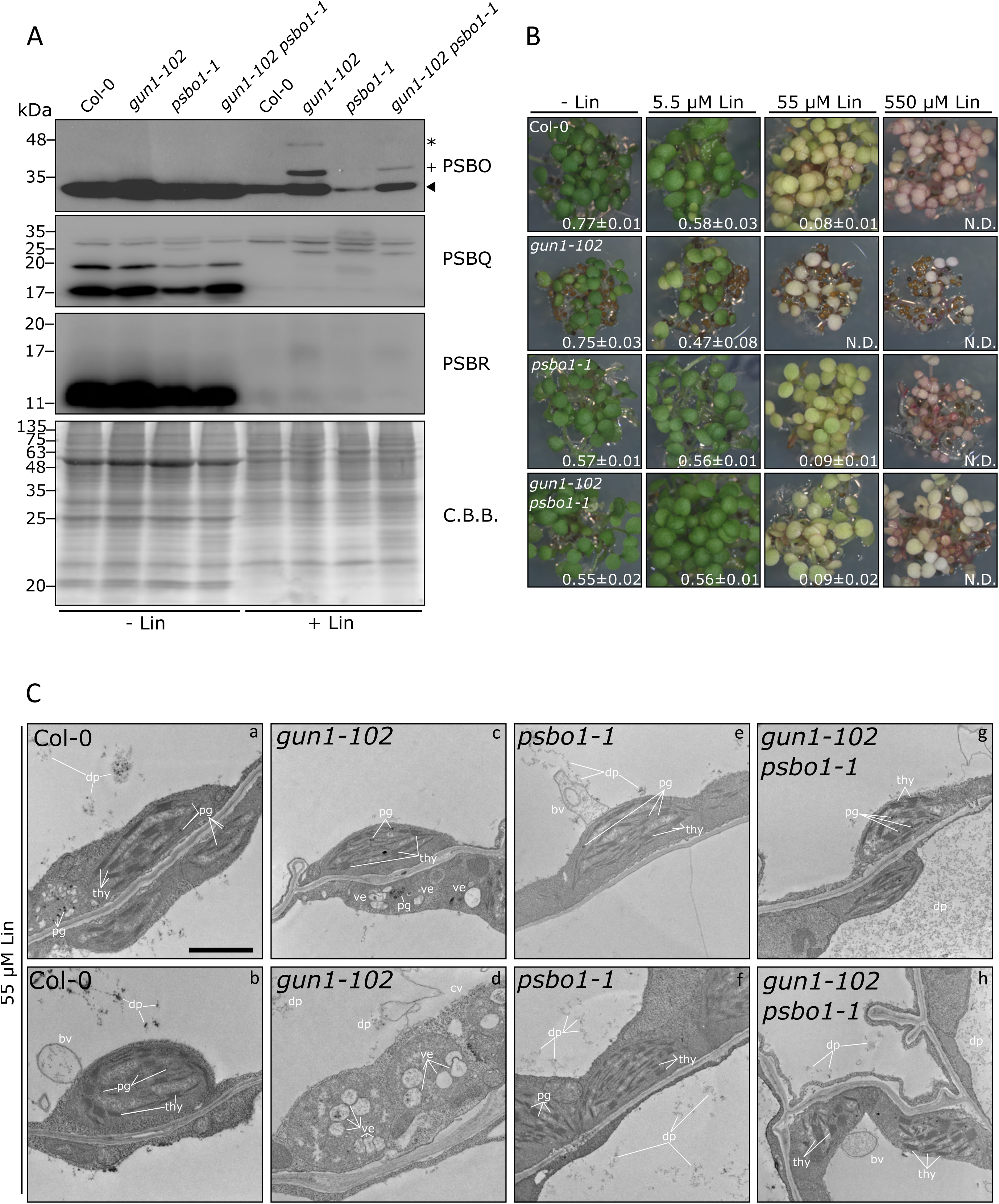
PsbO has a role in the increased sensitivity of *gun1* seedlings to lincomycin. **A**) Immunoblot analyses on Col-0, *gun1-102*, *psbo1-1* and *gun1-102 psbo1-1* total protein extract from seedlings grown in absence or presence of 550 µM Lin (-/+Lin) probed with PsbO, PsbR, PsbQ, antibodies. Coomassie Brilliant Blue (C.B.B.) staining of SDS-PAGE is shown as loading control. Note that PsbO antibody is targeting PsbO1 and PsbO2 and PsbO2 is detectable in *psbo1* mutant genetic background. Solid arrowhead indicates the mature PsbO protein, the plus symbol (+) indicates the precursor protein and the asterisk (*) possibly highlights post-translational modifications, i.e. ubiquitination. **B**) Visible phenotypes of 6 DAS wild-type (Col-0), single (*gun1-102*, *psbo1-1*) and double (*gun1-102 psbo1-1*) mutant seedlings grown on MS medium in the absence and presence of Lin in the growth medium at different concentration (5.5 µM, 55 µM, 550 µM). The Maximum Quantum Yield of Photosystem II (*Fv/Fm*) is reported as an indicator of photosynthesis efficiency and the functional state of chloroplasts (average ± SD; n ≥ 15). **C**) Transmission Electron Microscopy micrographs of mesophyll cells from 6 DAS seedlings of the indicated genotypes grown on MS medium with 55 µM Lin. Legend: thy) thylakoids; pg) plastoglobuli; ve) vesicle; bv) bugging vesicles; dp) degradation products; cv) collapsing vacuole. The scale bar represents 1 µm.

To investigate further the functional interaction between GUN1 and PsbO1 in the context of Lin-triggered plastid perturbation, a double *gun1-102 psbo1-1* mutant was generated and characterized. Immunoblot analysis performed on total protein extracts from Col-0, *gun1-102*, *psbo1-1* and *gun1-102 psbo1-1* seedlings grown in the absence or presence of 550 µM Lin, showed that PsbQ and PsbR subunits of OEC did not accumulate in mature forms in the different genetic background with Lin in the growth medium (**Figure 7 A**, **Figure S5 B**). On the contrary, PsbO proteins (PsbO1 and PsbO2) retained a relatively high accumulation of their mature form in all the genetic backgrounds grown in presence of Lin, particularly in all samples carrying the *gun1-102* mutation (**Figure 7 A**). In addition, PsbO signals were detected in two distinct bands at higher molecular weight in *gun1-102*+Lin samples (**Figure 7 A**), indicating the presence of precursor proteins retaining the chloroplast and thylakoid Transit Peptide (cTP and tTP, respectively), as shown by mass spectrometry analyses (**Table S6**), and possibly post-translationally modified PsbO proteins, likely due to poly-ubiquitination (**Figure 7A**, see asterisk). Moreover, all transcripts coding for OEC subunits (PsbO1, PsbO2, PsbQ1, PsbQ2, PsbP1, PsbP2 and PsbR) followed the trend of *Photosynthesis-Associated Nuclear Genes* (*PhANGs*), i.e. their expression was down-regulated by the presence of Lin in Col-0 and *psbo1-1* mutant, while they were relatively high in all the genetic backgrounds containing the *gun1-102* allele (**Figure S5 C**). These findings indicate that, unlike other photosynthesis-associated nuclear-encoded proteins, PsbO1 and PsbO2 can accumulate as mature proteins in non-green plastids and are not subjected to cytosolic degradation or post-transcriptional inhibition.

Since *gun1* shows enhanced sensitivity to Lin (Tadini *et al*, 2020c; Zhao *et al*, 2018), the double *gun1-102 psbo1-1* mutant was tested at increasing Lin concentrations (**Figure 7 B**). While Col-0 showed a significant decrease in greening and PSII photosynthetic performance at 55 µM Lin, the *gun1-102* mutant displayed no photosynthetic activity and no chloroplast accumulation at the same Lin concentration. Moreover, *gun1* seedlings showed a significantly lower *Fv/Fm* value at 5.5 µM Lin, corroborating the notion of a higher sensitivity to Lin, as reported previously (Tadini *et al*, 2020c; Zhao *et al*, 2018). On the other hand, the *psbo1-1* mutant showed little to no consequences when grown at 5.5 µM Lin in comparison to the -Lin control condition, behaving similarly to Col-0 at higher Lin concentrations. Strikingly, the introgression of *psbo1-1* mutation in *gun1-102* genetic background abolished the increased sensitivity to Lin as shown by the Col-0-like *Fv/Fm* values and the visible phenotype of *gun1-102 psbo1-1* seedlings grown on medium with 5.5 and 55 µM Lin (**Figure 7 B**). On the other hand, all genetic backgrounds showed no photosynthetic activity at 550 µM Lin.

To further investigate at plastid morphological level the partial restoration of *gun1-102* phenotype upon introgression of *psbo1-1* mutation, 6 DAS Col-0, *gun1-102*, *psbo1-1* and *gun1-102 psbo1-1* seedlings grown on medium with 55 µM Lin were observed through Transmission Electronic Microscopy (TEM) (**Figure 7 C**). Col-0 chloroplasts generally showed a lens-shaped appearance with intact thylakoid membranes. However, as expected from impaired photosynthetic parameters, alterations in chloroplast morphology were also observed. These include round-shape chloroplasts characterized by the presence of numerous plastoglobuli, together with membrane degradation products in the vacuole, and vesicles budding from the chloroplast envelope to the central vacuole (**Figure 7 C**, **panels a, b**). On the contrary, in *gun1-102* mesophyll cells, only a few thylakoid-containing chloroplasts were observed, while most of the plastids showed no properly formed thylakoid membranes, together with large plastoglobuli and vesicles filled with electron-dense material accumulating in the stroma (**Figure 7 C, panels c, d**). In a fraction of *gun1-102* cells, the vacuolar membrane appeared disrupted, suggesting ongoing vacuolar-mediated cell death (**Figure 7 C, panel d**). Intriguingly, *psbo1-1* mesophyll cells showed fully developed chloroplasts with large amounts of thylakoid membranes and little or no sign of perturbation (**Figure 7 C, panels e, f**), when compared to Col-0, indicating further that PsbO1-devoid plastids are less sensitive to chloroplast impairment and degradation when exposed to Lin. In accordance with this, the introgression of *psbo1-1* mutation in *gun1-102* genetic background led to a partial recovery of plastid/chloroplast impairment, visible in *gun1-102* samples. In particular, *gun1-102 psbo1-1* mesophyll cells were largely characterized by chloroplasts with properly shaped thylakoid membranes, while no severely damaged/degraded chloroplasts were found (**Figure 7 C, panels g, h**). Overall, these data indicate that PsbO, unlike any other Photosynthesis-related protein, accumulates in non-photosynthetic plastids with the scope of promoting their degradation upon impairment of chloroplast biogenesis.

### 2.6 PsbO involvement in chloroplast biogenesis and plastid quality control

The reduced accumulation of PsbO protein in *gun1-102 psbo1-1* mutant background correlates with its decreased sensitivity to Lin, measured as PSII photosynthetic efficiency, and with enhanced chloroplast integrity upon moderate (55µM) Lin-treatment (**Figure 7**). As PsbO proteins were found to physically interact with Chloroplast Vesiculation (CV), a protein involved in vesicle-mediated chloroplast degradation that plays a key role in chloroplast dismantling (Wang & Blumwald, 2014), and found in CV-induced vesicles, the role of PsbO1 superhub in chloroplast quality control was investigated. To this purpose, two viable independent *oePsbO1-GFP* lines were generated and isolated. At 12 DAS, both lines showed a pale variegated phenotype, with slightly reduced *Fv/Fm* values, when compared to Col-0 (**Figure 8 A**). A markedly lower *Fv/Fm* value and reduced growth rate were shown by *psbo1-1* knock-out line (Suorsa *et al*, 2016). The observed phenotype was confirmed at 18 DAS (**Figure S6 A**). The presence of the PsbO1-GFP protein was verified by immunoblotting (**Figure 8 B**). The accumulation of the endogenous PsbO protein was affected by the presence of PsbO1-GFP chimaeras causing slightly reduced accumulation. The subcellular localization of PsbO-GFP was analyzed in both lines by confocal microscope observations of protoplasts obtained from well-expanded leaves at 18 DAS. The GFP signal appeared both as dense foci and diffused fluorescence within green plastids co-localizing with chlorophyll fluorescence, but it was also detected both in vesicle-like structures budding from chloroplasts and non-overlapping with chlorophyll autofluorescence and as independent vesicles detached from the organelles (**Figure 8 C**). More specifically, PsbO1-GFP large fluorescent foci were observed mostly in vesicles that did not overlap with chlorophyll autofluorescence and that showed TIC20-RFP accumulation, proving the plastid origin of such structures (**Figure S7**).

**Figure 8.**
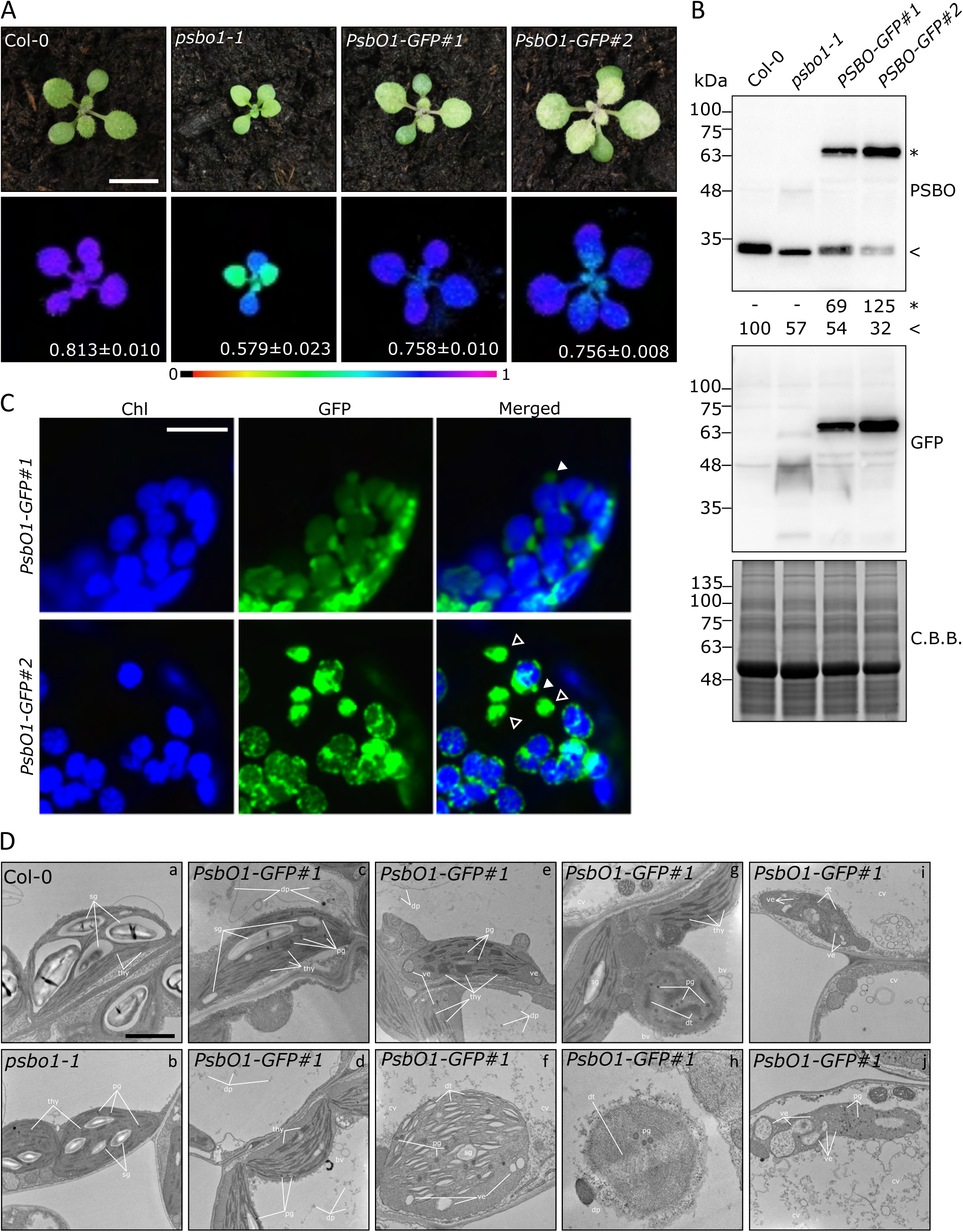
PsbO involvement in chloroplast quality control and degradation. **A**) Pictures of the visible phenotypes displayed by 12 DAS plantlets of the indicated genotypes and Imaging PAM pictures representing in false colours the *Fv/Fm* parameter and the average ± SD values (n ≥ 4). The scale bar is 1 cm. **B**) Immunoblot analyses on total protein extracts from Col-0, *psbo1-1* and the two independent *oePSBO1-GFP* lines grown on soil until 12 DAS probed with PsbO- and GFP-specific antibodies. Coomassie Brilliant Blue (C.B.B.) staining of SDS-PAGE is shown as loading control. Asterisk (*) and arrowhead (<) indicate the chimaera and the endogenous PsbO, respectively. Numbers indicate the PsbO signal intensities relative to Col-0. **C**) Fluorescence signals from chlorophylls (Chl, blue) and GFP (green) detected in protoplasts obtained from leaves of the indicated genotypes observed through the confocal microscope. Solid arrowheads indicate budding vesicle-like structures while empty arrowheads show detached vesicles. The scale bar is 10 µm. **D**) Transmission Electron Microscopy micrographs of mesophyll cells of Col-0 (a), *psbo1-1* (b) and *PsbO1-GFP* (c-j) lines from 12 DAS plantlets grown on soil until 12 DAS. Legend: thy) thylakoids; dt) degrading thylakoids; sg) starch granule; pg) plastoglobuli; ve) vesicle; bv) bugging vesicles; dp) degradation products; cv) collapsing vacuole. The scale bar represents 1 µm.

As PsbO was found in CV-induced degradation vesicles (Wang & Blumwald, 2014), the possibility of ongoing chloroplast degradation at the basis of *PsbO1-GFP* chlorotic/variegated phenotype was investigated by TEM analyses (**Figure 8 D**). Under physiological conditions, Col-0 leaves displayed chloroplasts with thylakoid structures organized in grana stacks and stroma lamellae, together with a large number of starch granules (**Figure 8 D, panel a**), while *psbo1-1* chloroplasts showed swollen thylakoid structures and increased amount of plastoglobuli, as OEC seems to play a role in the structural integrity of thylakoid membranes (Suorsa & Aro, 2007), together with a reduced amount of starch granules (**Figure 8 D, panel b**). On the other hand, chloroplasts obtained from *PsbO1-GFP* lines showed a wide range of alterations in plastid morphology in line with the variegated phenotype: from *psbo1*-like chloroplasts (**Figure 8 D, panels c, d**) to more severely impaired ones, with highly damaged and degenerated thylakoid membranes (**Figure 8 D, panels e, f, g**), to completely dismantled chloroplast-like organelles (**Figure 8 D, panels h, i, j**). More into details, several structures associated with chloroplast dismantling could be observed, as round-shaped chloroplasts detached from the plasma membrane migrating towards the vacuole, compatible with the micro-autophagy process (**Figure 8 D, panels g, h**; Zhuang and Jiang, 2019; Woodson, 2022), or highly vesiculated chloroplasts compatible with fission-type ATG- and PUB4-dependent micro- and macro-autophagy (**Figure 8 D, panels e, f, i, j** Jeran et al., 2021; Lemke et al., 2021; Woodson, 2022). Moreover, vesicles forming in chloroplasts and migrating towards the central vacuole, similar to CV-induced degrading vesicles, were observed (**Figure 8 D, panel d, e**; Wang and Blumwald, 2014). Chloroplast degradation led to the accumulation of electron-dense material in the central vacuole, together with large amounts of membranes and degradation products (**Figure 8 D, panels c-j),** resembling what previously observed in *gun1 ftsh2* and *gun1 ftsh5* genetic backgrounds (Tadini *et al*, 2020c). In several cells, chloroplast degradation was followed by vacuolar collapse and dismantling of the entire cell structure (**Figure 8 D, panel g, i, j**), similar to what was described in ROS-induced cell death (Woodson *et al*, 2015).

To investigate whether the observed variegated phenotype displayed by the *PsbO1-GFP* lines (**Figure 8**) could be linked to CV activity, the *oePsbO1-GFP#1* line was crossed with the *amiR-CV* line, in which *CV* gene expression is silenced by an artificial microRNA (Wang & Blumwald, 2014). To compare phenotypes, Col-0, *psbo1-1*, *oePsbO1-GFP#1*, *amiR-CV* and the double mutant *oePsbO1-GFP#1 amiR-CV* were cultivated on soil in controlled conditions (**Figure 9 A**). Strikingly, the obtained double mutant phenotype was partially restored towards a Col-0-like appearance. At 12 DAS, leaves were nearly devoid of variegations and photosynthetic parameters were slightly improved. At 18 DAS, the recovery of all parameters was even more appreciable (**Figure S6 B**). The accumulation of PsbO1-GFP chimaera was confirmed with immunoblots incubated with anti-PsbO and anti-GFP antibodies (**Figure S8**). Observed at the confocal microscope, protoplasts isolated from mesophyll tissues of *PsbO1-GFP#1 amiR-CV* showed the GFP signal as foci within chloroplasts or diffused matching the chlorophyll autofluorescence (**Figure 9 B**). Moreover, the signal observed in the double mutant was detected only within chloroplasts, whereas in the *oePsbO1-GFP#1* line, foci outside the plastids were detectable. These data support the idea that an imbalance of PsbO1 accumulation could be a trigger for CV-mediated chloroplast dismantling. In order to visualize such functional interaction, protoplasts obtained from the *PsbO1-GFP#1* line were transiently transformed with a construct expressing the *CV* coding sequence fused to *RFP* under the control of the *CaMV35S* promoter and observed by confocal microscope. As a control, *PsbO1-GFP#1* protoplasts were analyzed as well. Strikingly, the RFP signal resulted to be diffusing in the cytoplasm and as foci matching the GFP fluorescence, supporting the co-localization of PsbO aggregates inside CV-induced degradation vesicles (**Figure 9 C**).

**Figure 9.**
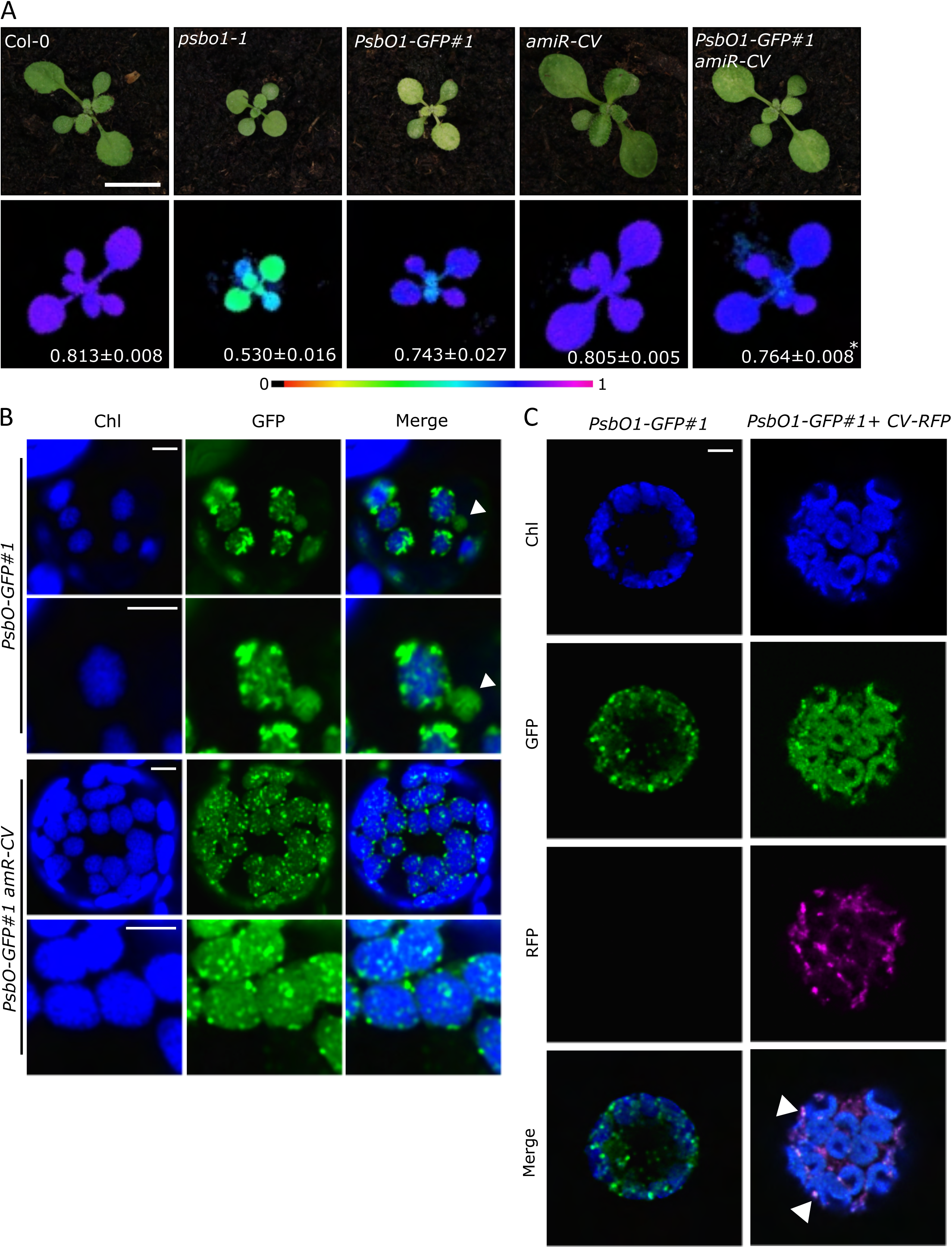
PsbO functional interaction with CV. **A**) Visible phenotypes of 12 DAS plantlets of the indicated genotypes and Imaging PAM pictures representing in false colours the *Fv/Fm* parameter and the average ± SD values (n ≥ 4). The scale bar represents 1 cm. **B**) Fluorescence signals from chlorophylls (Chl, blue) and GFP (green) detected in protoplasts obtained from leaves of the indicated genotypes observed through the confocal microscope at different magnifications. Solid arrowheads indicate budding vesicle-like structure. The scale bar represents 5 µm. **C**) Fluorescence signals from chlorophylls (Chl, blue), GFP (green) and RFP (magenta) detected in protoplasts obtained from *PsbO1-GFP#1* line and transformed with *CV-RFP* construct observed through the confocal microscope. The scale bar represents 5 µm.

## 3. Discussion

Intracellular communication pathways rely on a complex network of molecular interactions that exert control over organelle development and play a key role in adaptation to the environment (Ng *et al*, 2014; Wu & Bock, 2021). While GUN1 seems to play a central role in transmitting information from developing and mature chloroplasts across the cell with the aim to adjust the nuclear gene expression, its primary functional role is still under debate (Koussevitzky *et al*, 2007; Zhao *et al*, 2019b; Tadini *et al*, 2020b; Wu *et al*, 2019a; Tadini *et al*, 2020c; Colombo *et al*, 2016). Based on literature data, it is clear that Lin-treated samples show lost/reduced accumulation of most plastid-located proteins, as a consequence of i) inhibition of plastid translation, ii) gene expression down-regulation of nuclear-encoded and plastid-located subunits, iii) post-transcriptional suppression of plastid-located protein accumulation at cytosolic level, by either translation inhibition or degradation (Koussevitzky *et al*, 2007; Tadini *et al*, 2020c; Wu *et al*, 2019b). To shed further light on this matter, we attempted the holistic description of the Arabidopsis seedling proteome in response to the inhibition of plastid translation and differentiation into chloroplasts (by Lin), in a genetic context devoid of GUN1-dependent plastid-to-nucleus signaling pathway.

### GUN1 coordinates a stress adaptive response that involves plastids and extra-plastid cell compartments

Upon impairment of plastid translation, Photosynthesis-related proteins are generally suppressed, whereas few plastid-located subunits are retained (e.g. ClpC1, ClpC2, CPN60s), and they are responsible for the reorchestration of plastid functions (**Figure 2**; **Figure 5; Figure S2**; **Table S1**). In this context, plastid-localized chaperones, proteases and ROS scavengers showed increased topological relevance as hubs only in Col-0+Lin network models (**Figure 10 a**), indicating that, upon Lin-treatment, housekeeper and stress-response plastid functions are promoted in a GUN1-dependent manner. Accordingly, GUN1 was found to physically interact with the plastid protein homeostasis machinery, including chaperones and proteases such as ClpC2 (Tadini *et al*, 2016; Wu *et al*, 2019a; Jia *et al*, 2019). This is also in line with the enhanced accumulation of GUN1 protein itself upon impairment of plastid functions and correlates with the increased sensitivity of *gun1* seedlings to heat, cold, chemicals (Lin and Norflurazon) and with the albino/variegated phenotypes shown when *gun1* is combined with mutants altered in plastid protein homeostasis (**Figure 7**; Tadini, Peracchio, et al., 2020; Marino et al., 2019; Lasorella et al., 2022; Zhao et al., 2018; Tadini, Jeran and Pesaresi, 2020; Tadini, Jeran, Peracchio, et al., 2020). Overall, these pieces of evidence corroborate the notion that GUN1 is a key factor in plastid stress response, integrating multiple adaptive mechanisms (**Figure 10 a**).

**Figure 10.**
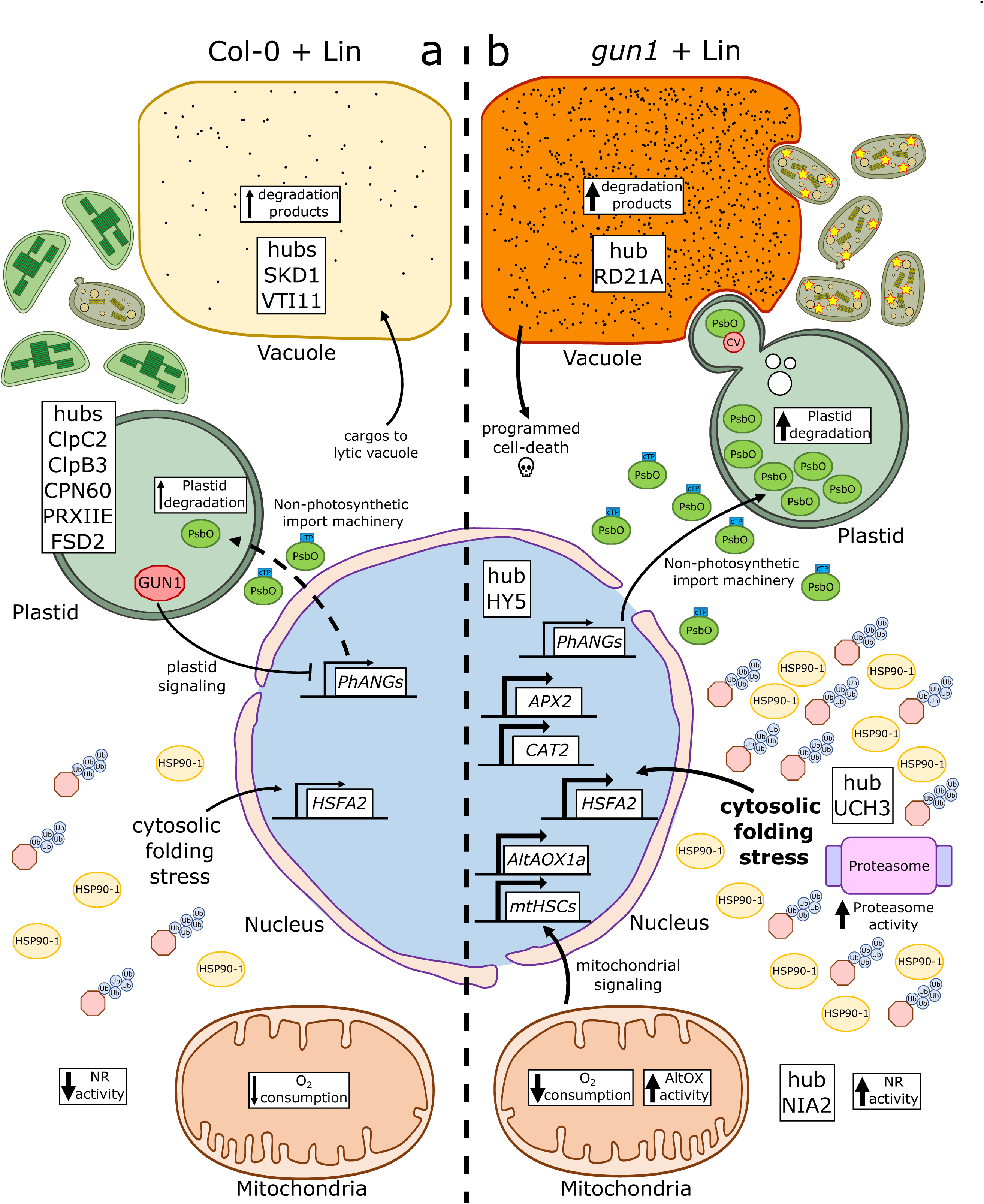
Working model of main protein hubs and cellular events described in presence (a) or absence of GUN1 (b) upon chloroplast biogenesis impairment due to the presence of Lin in the growth medium. **a)** GUN1 promotes the reorchestration of plastid proteome to overcome the Lin-mediated protein synthesis inhibition. This plastid-located response involves non-photosynthetic proteins, such as chaperones and stress-related proteins (ClpC2, ClpB3, CPN60, PRXIIE, FSD2) and promotes the repairment of developing chloroplasts. In this context, plastid translation inhibition lowers NR activity and O_2_ consumption in mitochondria and elevates to hubs the vacuole-related proteins SKD1 and VTI11. **b)** In the absence of GUN1, plastid-located compensatory response and plastid-to-nucleus signalling fail while developing chloroplasts become more sensitive to stress challenges, including presence of Lin. Consequently, plastid degradation dependent on vacuole activity is enhanced by activation of micro- and macroautophagy pathways. Eventually, the vacuole-mediated cell death is triggered, as evidenced by the RD21A vacuolar hub and the observed dissolving tonoplast. At the same time, the abolished GUN1-mediated repression of *PhANGs*-encoded proteins leads to higher cytosolic folding stress consisting in increased accumulation of precursor proteins. The cytosolic compartment compensates by enhancing cytosol detoxification activity, elevating NIA2 and UCH3 as hubs, and promoting NR activity, chaperone accumulation, protein ubiquitination level and proteasome activity. In line with the fact that the nuclear gene expression differs in the two genetic backgrounds, the HY5 transcription factor, related to photomorphogenesis, was found as a hub in *gun1*+Lin conditions. Mitochondria adapt as well to the exacerbated stress by increasing their folding capacity and stimulating the AltOX activity. Intriguingly, PsbO1 was the unique plastid-located hub in *gun1*+Lin, pointing to a role in plastid quality assessment, besides its involvement in photosynthesis.

Furthermore, whole-cell reprograming and reorchestration of plastid functions following Lin treatment are accompanied by a robust enrichment in GO terms associated with nuclear activities, Golgi vesiculation and mitochondrial functional reorganization. As a matter of fact, the GUN1-mediated repression of *Photosynthesis-Associated Nuclear Genes* (*PhANGs*) occurring in Col-0 samples treated with Lin has been indicated to derive from the enhanced cytosolic folding stress and implies cytosolic degradation of *PhANGs*-related transcription factors (Wu *et al*, 2019a; Tadini *et al*, 2020b, 2020a). Indeed, our analyses identified an increase in the abundance of cytosolic stress-related proteins and transcripts in Col-0+Lin samples, together with higher protein ubiquitination (**Figures 4-7**; Tadini, Peracchio, et al., 2020; Jeran et al., 2021). Network topological analysis also evidenced that many vacuole-related proteins acquire the hub status when Lin is applied in both Col-0 and *gun1* backgrounds, remarking the role of the vacuole in the clearance of protein aggregates and defective organelles (Muntz, 2007; Woodson, 2022; Shimada *et al*, 2018; Tan *et al*, 2019). Moreover, the vacuolar proteins SKD1 and VTI11, both involved in vacuolar protein trafficking and delivery of protein cargos to lytic vacuoles (Muntz, 2007; Sanmartín *et al*, 2007; Shahriari *et al*, 2010), were among the main topological players in Col-0+Lin condition.

### Plant cells devoid of GUN1 rely on extra-plastid compartments to address stress challenges

Strikingly all the plastid hubs identified in the proteomes of Col-0+Lin seedlings lost any relevance in *gun1*+Lin samples. Conversely, the role of hubs in *gun1*+Lin was assumed by non-plastid proteins (**Figure 10 b**). Among them, UCH3 stands out as the cytosolic hub involved in protein ubiquitination (**Figure 5 A**), as a consequence of the abolished GUN1-mediated *PhANGs* suppression and the increased cytosolic folding stress observed in *gun1*+Lin seedlings with respect to Col-0+Lin (Tadini *et al*, 2020c). In agreement with this finding, protein ubiquitination was enhanced in *gun1*+Lin samples, together with the accumulation of chaperones and proteasome activity (**Figure 6 A, B**). As a consequence, the nuclear gene expression was modulated accordingly (**Figure 6 C**). Intriguingly, the HY5 transcription factor, which drives photomorphogenesis and light-responsive gene expression (Zhao *et al*, 2019a), was identified as the PPI hub only in *gun1+*Lin condition. Furthermore, the RD21A protease, a protein released by the vacuole during the programmed cell death cascade (Koh *et al*, 2016; Boex-Fontvieille *et al*, 2015; Wang *et al*, 2008), was elevated as a hub in *gun1+*Lin condition, in place of SKD1 and VTI11 observed in Col-0+Lin. It is tempted to speculate that upon failure of GUN1-mediated plastid response, the vacuole compartment is capable of initiating the programmed cell death, as supported by the presence of highly disrupted plastids together with degenerated tonoplasts in Lin-treated seedlings (**Figure 7**; Hatsugai et al., 2004), and well correlates with the altered redox state of *gun1*+Lin, as previously described (Fortunato *et al*, 2022). This status was further evidenced by NIA2 as PPI hubs in *gun1+*Lin sample, together with the higher accumulation of cytosolic-located ROS/RNS scavenging factors and the increased NR activity (**Figure 6**). Additionally, mitochondria were found to play an important role in the cellular events triggered by the Lin treatment as indicated by the PPI functional modules (the mitochondrial respirasome complex) and the GO term analysis, in which the Organelle Organization and Metabolism categories appeared enriched (**Figure 2**; **Figure 5 and Figure S2**). In line with this, the activation of mitochondrial alternative oxidase and mitochondrial UPR gene expression appeared to be specific for *gun1-102+*Lin responses (**Figure 6**). Noteworthy, AltOX is an important marker of active mitochondrial retrograde signaling and a variety of studies have shown that its expression is increased when chloroplasts are hampered (Koussevitzky *et al*, 2007; Giraud *et al*, 2009), supporting further the compensatory role of mitochondria in cellular adaptive responses when plastids are unable to cope with stress (**Figure 10 b**).

### PsbO subunit has an atypical behavior with respect to photosynthesis related proteins and promotes plastid degradation when mis-located in plastid stroma

Following the holistic evaluation of the proteome profiles, graph theory-based computational strategies surprisingly depicted PsbO1 as a superhub, being a PPI hub in both genotypes in the control condition (-Lin) and the only plastid-located co-expression hub in *gun1*+Lin samples (**Figure 7**). PsbO, encoded by a two-copy gene, is involved in the linear electron transport across the thylakoid membrane, being a subunit of oxygen-evolving complex (OEC), responsible for water splitting and P680 regeneration at lumenal side of PSII (Allahverdiyeva *et al*, 2013; Yi *et al*, 2005). Almost the entirety of *PhANGs*-encoded proteins does not accumulate upon Lin treatment, while the related transcripts are suppressed by retrograde signaling pathways and a minor fraction is regulated at the post-transcriptional level (Wu *et al*, 2019b). In the absence of GUN1, *PhANGs* down-regulation fails and protein accumulation is prevented by translation inhibition and degradation at the cytosolic level, although cytosolic precursor proteins can be observed in *gun1-102*+Lin samples (**Figure 7**; Tadini, Peracchio, et al., 2020; G.,-Z., Wu et al., 2019; Jeran et al., 2021). Strikingly, the accumulation of PsbO protein was detected in plastids devoid of photosynthetic machinery and thylakoid membranes, as in the case of *tic56* and *ppi2* Arabidopsis mutants or Spectinomycin- and Lin-treated seedlings (Köhler *et al*, 2016). Moreover, PsbO accumulates as a mature protein out of stoichiometry when compared to other OEC and PSII subunits, suggesting an independent role besides OEC-related functions, possibly in chloroplast quality assessment and degradation. The lack of GUN1 caused PsbO to accumulate to a higher level in plastids treated with Lin and its decreased abundance directly correlates with the reduced sensitivity of seedlings to Lin, as observed in *gun1-102 psbo1-1* compared to *gun1-102* samples (**Figure 7**). In previous studies, PsbO was identified in Chloroplast Vesiculation-induced bodies and found to be involved in protein-protein interaction with CV itself (Wang & Blumwald, 2014). Accordingly, the over-accumulation of PsbO1-GFP protein correlated with chloroplast degradation, partially recovered upon *CV* down-regulation (**Figure 8, 9**; see also Woodson, 2022; Wang and Blumwald, 2014). Furthermore, budding vesicles observed in *PsbO1-GFP* chloroplasts strongly resemble micro- and macro-autophagy processes (Jeran *et al*, 2021; Lemke & Woodson, 2022; Lemke *et al*, 2021), while no vesicle was detected upon *CV* downregulation. PsbO1-GFP accumulation also resulted in vacuolated structures and disrupted vacuolar membrane resembling those observed in cells subjected to programmed cell death (Woodson *et al*, 2015; Tadini *et al*, 2023). This well correlates with the identification of vacuolar-related hubs, including RD21A (Koh *et al*, 2016), among *gun1*+Lin co-expression hubs. It has been suggested that, rather than the over-accumulation of PsbO itself, the altered localization of PsbO-GFP in the stromal compartment is the source of the variegated/chlorotic phenotype (Jiang *et al*, 2017). In this context, the simultaneous impairment of thylakoid biogenesis, together with the non-downregulation of *PsbO* gene expression, would cause the accumulation of PsbO protein in the stromal compartment and lead to enhanced chloroplast degradation in *gun1*+Lin seedlings (**Figure 10 b**).

## 4. Conclusions and future perspectives

Through a system biology approach, we have provided new insights into the molecular mechanisms activated by the inhibition of chloroplast biogenesis in the presence/absence of the GUN1-mediated plastid-to-nucleus signaling pathway. Along with the description of proteomic rearrangements and the identification of proteins that assume the role of network hubs, we provide evidence of adaptive responses triggered by plastid protein mis-localization. The accumulation of *PhANGs*-encoded proteins in the cytosol is known to activate a cytosolic folding stress response, responsible for a feedback loop control aimed to inhibit translation and/or promote pre-protein degradation (Tadini *et al*, 2020c; Wu *et al*, 2019b), followed by the activation of vacuole- and mitochondria-located compensation mechanisms. On the other hand, the mis-localization of PsbO in the plastid stromal compartment appears to act as a signal for the degradation of altered plastids with the aim of preventing ROS formation and oxidative damage. Overall, these data highlight the importance of plastid protein mis-localization as a source of adaptive responses to different chloroplast development stages and environmental conditions, aimed at repairing or dismantling damaged plastids.

## 5. Materials and Methods

### Plant material and growth conditions

*Arabidopsis thaliana* wild-type (Col-0) and mutant plants were grown on soil in long-day controlled conditions (150 μmol m^−2^ sec^−1^ 16 h/8 h light/dark cycles). *gun1-102, psbo1-1* and *psbq1-1 psbq2-1 psbr-1* T-DNA insertional mutants were previously described (Lundin *et al*, 2007; Tadini *et al*, 2016; Allahverdiyeva *et al*, 2013). *PsbO1-GFP* lines were obtained by Agrobacterium-mediated transformation of Col-0 plants with *Agrobacterium tumefaciens* strain transformed with the pB2GW7 plasmid (gatewayvectors.vib.be), carrying the coding sequences of *PsbO1* gene under the control of CaMV35S promoter and cloned in frame with the *GFP* coding gene. *amiR-CV* line was previously described (Wang & Blumwald, 2014). *PsbO1-GFP amiR-CV, PsbO1-GFP TIC20-RFP* and *gun1-102 psbo1-1* double mutants were obtained by manual crossing and segregation analysis. For the analysis of mitochondria morphology and dynamics, *gun1-102* plants were crossed with *Arabidopsis thaliana* Col-0 lines expressing the Yellow Fluorescent Protein (YFP) at the mitochondrial level, available in the laboratory of Prof. Michela Zottini. For Lin treatment, seeds were surface-sterilized and grown for 6 days (80 μmol m^−2^ sec^−1^ on a 16 h/8 h dark/light cycle) on Murashige and Skoog medium (Duchefa) supplemented with 2% (w/v) sucrose, 1.5% (w/v) Phyto-Agar (Duchefa) and Lin at 550 µM, unless differently indicated. Primers for mutant isolation and cloning are listed in **Table S7**.

### LC-MS/MS analyses

For proteomic analysis, 6 DAS seedlings grown on Murashige and Skoog medium supplemented with 2% (w/v) sucrose, 1.5% Phyto-Agar and 550 µM Lin were homogenized in SDS lysis buffer [4% (w/v) SDS, 100 mM Tris/HCl pH 7.5, 100 mM DTT], incubated at 95°C for 5 min and centrifuged at 13.000 g for 5 min. Crude protein extracts were digested by trypsin using two digestion FASP methods (Wiśniewski, 2018). Prior to mass spectrometric analysis, peptides were dried in a speed-vac and desalted using Zip-Tips (lC18; Millipore). 4 µg of peptides were analyzed in a QExactive mass spectrometer coupled to a nano EasyLC 1000 (Thermo Fisher Scientific). Solvent composition at the two channels was 0.1% (v/v) formic acid for channel A and 0.1% (v/v) formic acid, 99.9% (v/v) acetonitrile for channel B. 4µL of each sample were loaded on a self-made column (75 µm×150 mm) packed with reverse-phase C18 material (ReproSil-Pur 120 C18-AQ, 1.9 µm, Dr. Maisch GmbH) and eluted at a flow rate of 300 nL/min by a gradient from 2 to 35% (v/v) B in 80 min, 47% (v/v) B in 4 min and 98% (v/v) B in 4 min. Samples were acquired in a randomized order. The mass spectrometer operated in data-dependent mode (DDA), acquiring full-scan MS spectra (300-1700 m/z) at a resolution of 70000 at 200 m/z after accumulation to a target value of 3000000, followed by HCD (higher-energy collision dissociation) fragmentation on the twelve most intense signals per cycle. HCD spectra were acquired at 35000 resolution, using a normalized collision energy of 25 and a maximum injection time of 120 ms. The automatic gain control (AGC) was set to 50000 ions. Charge state screening was enabled and single and unassigned charge states were rejected. Only precursors with an intensity above 8300 were selected for MS/MS (2% underfill ratio). Precursor masses previously selected for MS/MS measurement were excluded from further selection for 30s, and the exclusion window was set at 10 ppm. Samples were acquired using internal lock mass calibration on m/z 371.1010 and 445.1200. For identification and characterisation of PsbO proteins, following Western Blot analysis the corresponding protein bands were in-gel digested with trypsin and analysed by quadrupole ion trap mass spectrometry as described previously (Domingo *et al*, 2023). PsbO proteins were identified using Turbo SEQUEST Bioworks™ 3.3 software (Thermo Electron Corporation, California, USA) against the *A. thaliana* database (Araport11, www.arabidopsis.org/download, downloaded 1 December 2023) as described elsewhere (Domingo *et al*, 2023).

### Raw data processing

Raw data, from literature and newly generated, were processed by the Sequest HT algorithm in Proteome Discoverer 2.1 software (Thermo Fisher Scientific). Experimental MS/MS spectra were compared with theoretical spectra obtained by in silico digestion of 39345 protein sequences downloaded from UNIPROT, in June 2020 (www.uniprot.org). Criteria of searching were set as: trypsin enzyme, three missed cleavages per peptide, mass tolerances of ± 50 ppm for precursor ions and ± 0.8 Da for fragment ions. The percolator node was used with a target-decoy strategy to give a final false discovery rate (FDR) ≤0.01 (strict) based on q-values, considering a maximum deltaCN of 0.05. Only peptides with a minimum length of 6 amino acids, confidence at a “High” level and rank 1 were considered. Protein grouping and strict parsimony principle were applied.

### Proteomics datasets

Proteomics data used for this study were collected from three independent mass-spectrometry experiments. The mass spectrometry proteomics data generated in this study (Dataset 1, total protein content from Col-0±Lin and *gun-102*±Lin 6 days old seedlings) has been deposited to the ProteomeXchange Consortium via the PRIDE (Perez-Riverol *et al*, 2022) partner repository with the dataset identifier PXD051970. Dataset 2 originated from soluble proteins (Col-0±Lin; *gun-102*±Lin, 6 days old seedlings) and was previously published (Tadini *et al*, 2020c). Dataset 3 originated from total protein content (Col-0±Lin; *gun-101*±Lin, 5 days old seedlings) (Wu *et al*, 2019a).

### Gene ontologies in protein profiles

A preliminary functional evaluation of protein profiles was performed by the GO Term Enrichment for Plants tool in TAIR (Berardini *et al*, 2015) via Panther (Mi *et al*, 2021) database. Using the protein profile of each biological replicate, enriched Biological Processes (BPs) and Molecular Functions (MFs) were retrieved (*P* ≤ 0.05). Fisher’s Exact test and Bonferroni correction for multiple testing were set. Enriched BPs and MFs were compared by LDA and those with F ratio ≥ 5 and P-value ≤ 0.01 were retained.

### Label-free quantitation based on spectral counting

The spectral count (SpC) values of the identified proteins were normalized using a total signal normalization method and compared using a label-free quantification approach (Vigani *et al*, 2017). Specifically, data matrix dimensionality (n total = 36; n=9 per condition) was reduced by LDA. Pairwise comparisons (Col-0 vs Col-0+Lin; Col-0 vs *gun1*; *gun1* vs *gun1*+Lin; Col-0+Lin vs *gun1*+Lin) were performed and only proteins with F ratio ≥ 5 and P-value ≤ 0.01 were considered. Fold change was estimated using the natural logarithm of the average spectral count (avSpC) ratio. The fold change value of proteins identified exclusively in one of the two compared conditions was set to ±5. Proteins selected by LDA were processed by Principal Component Analysis and Hierarchical Clustering applying Ward’s method and Euclidean distance metric. Data processing was performed via JMP 15.1 SAS software.

### Reconstruction of functional PPI network modules differentially expressed

*Arabidopsis thaliana* PPI network model was reconstructed from the higher-confident differentially abundant proteins (DAPs, n=326, P≤0.001) found comparing the characterized proteome profiles (Col-0 vs Col-0+Lin; Col-0 vs *gun1*; *gun1* vs *gun1*+Lin; Col-0+Lin vs *gun1*+Lin). To reconstruct the networks, StringApp (Doncheva *et al*, 2019) of Cytoscape (Su *et al*, 2014) was used. Protein-protein interactions were filtered by retaining exclusively those annotated in databases and/or experimentally determined with a Score ≥0.15. Proteins were grouped in functional modules by the support of BINGO 2.44 (Maere *et al*, 2005); *Arabidopsis thaliana* organism, hypergeometric test, Benjamini–Hochberg FDR correction and a significance level ≤0.01 were set. Reconstructed networks were visualized with Cytoscape; node color code indicates up-regulated (red) and down-regulated (light blue) proteins based on average Spectral count (avSpC) normalization (SpC normalized in the range 0-100).

### Protein-protein interaction and co-expression network models

Arabidopsis PPI network models were reconstructed from all proteins identified in different conditions (Col-0, *gun1*, Col-0+Lin, *gun1*+Lin). To reconstruct the networks, StringApp (Doncheva *et al*, 2019) of Cytoscape (Su *et al*, 2014) was used. Retrieved protein-protein interactions were filtered by retaining exclusively those annotated in databases and/or experimentally determined, with a Score ≥0.6. According to these parameters, a fully connected network of 4669 nodes and 111438 edges was built. Protein co-expression network models were reconstructed by processing the Arabidopsis protein profiles in each condition (Col-0, *gun1*, Col-0+Lin, *gun1*+Lin). To evaluate how correlation change among the considered conditions, Spearman’s rank correlation coefficient was computed only for proteins (n = 837) identified in all analyzed samples (n = 36, 9 per condition); Spearman’s rank correlation score ≥ |0.95| and P ≤ 0.01 were set as thresholds. Correlation network models were processed with Cytoscape.

### Topological evaluation of PPI and co-expression network models

Topological network analysis was performed with Cytoscape’s plugin NetworkAnalyzer and Centiscape 2.2 (Assenov *et al*, 2008; Scardoni *et al*, 2015), as previously reported (Di Silvestre *et al*, 2021). Betweenness, Centroid and Bridging centralities were calculated for PPI models, while node Degree was calculated for co-expression ones. For both models, nodes with centrality values above the average calculated on the whole network were considered hubs (Di Silvestre *et al*, 2018). The statistical significance of topological results was tested by considering randomized network models; they were reconstructed and analyzed with an in-house R script based on VertexSort (to build random models), igraph (to compute centralities), and ggplot2 (to plot results) libraries.

### Protein sample preparation and immunoblot analyses

Immunoblots were performed on total protein extracts. Fresh plant material was homogenized in sample buffer containing 20% (v/v) glycerol, 4% (w/v) SDS, 160 mm Tris–HCl pH 6.8, 10% (v/v) 2-mercaptoethanol (10 µL of buffer every mg of plant material; (Tadini *et al*, 2023). Samples were incubated at 65 °C for 15 min and centrifuged for 10 min at 16 000 g to pellet debris. Cleared homogenates were incubated for 5 min at 95 °C. 40 µL of protein samples, corresponding to 4 mg of fresh weight, were fractionated by SDS–PAGE 10% (w/v) polyacrylamide (Schägger & von Jagow, 1987) and then blotted to polyvinylidene difluoride (PVDF) membranes (0.45 µm pore size). Filters were immunodecorated with specific antibodies. Antibodies against PsbO (AS05 092) PsbQ (AS06 142-16), PsbR (AS05 059), UBQ11 (AS08 307A) and HSP90-1 (AS08 346) were obtained from Agrisera (www.agrisera.com/), GFP antibody (A-6455) was obtained from Invitrogen (www.thermofisher.com) and anti-RFP was purchased from Chromotek. Signal detection was performed by ChemiDoc Touch (Bio-Rad; www.bio-rad.com) and Image Lab software (Bio-Rad; www.bio-rad.com).

### Transmission electron microscopy (TEM)

TEM analyses were performed as described previously (Jeran *et al*, 2021). Plant material was vacuum-infiltrated for 4 h in a buffer containing 2.5% (v/v) glutaraldehyde and 100 mM sodium cacodylate and incubated overnight at 4°C. Samples were post-fixed in 100 mM cacodylate solution containing 1% (w/v) osmium tetroxide for 2 h at 4 °C. Then, samples were incubated overnight at 4 °C in the dark with 0.5% (w/v) uranyl acetate to carry out counterstaining. Tissue dehydration was achieved by 10 min incubation in ethanol of increasing concentrations (70%, 80%, 90%; v/v). Samples were incubated in 100% ethanol for 15 min and then in 100% propylene oxide for 15 min, twice. Epon-Araldite resin was obtained by mixing Embed-812, Araldite 502, dodecenylsuccinic anhydride (DDSA) and Epon Accelerator DMP-30. Samples were infiltrated for 2 h with a 1:2 mixture of Epon-Araldite and propylene oxide, then infiltrated for 1 h with a 1:1 mixture of Epon-Araldite and propylene oxide and finally incubated overnight at room temperature in a 2:1 mixture of Epon-Araldite and propylene oxide. Samples were embedded in pure resin before polymerisation at 60°C for 48 h. Ultra-thin sections of 70 nm were obtained by a diamond knife (Ultra 45°, DIATOME) and placed on copper grids (G300-Cu, Electron Microscopy Sciences). Sample observation was achieved by transmission electron microscopy (Talos L120C, Thermo Fisher Scientific) at 120 kV and images were collected with a digital camera (Ceta CMOS Camera, Thermo Fisher Scientific).

### Proteasome activity

Proteasome activity was determined according to previous work (Paradiso *et al*, 2020). Arabidopsis seedlings (0.1 g) were ground in liquid nitrogen, homogenized in a 1:3 (w:v) ratio with extraction buffer (50 mM Hepes-KOH, pH 7.2, 2 mM DTT, 2 mM ATP and 250 mM sucrose) and centrifuged at 20000g for 15 min at 4°C. Ten microliters of supernatants, with 1 mg/ml protein concentration, were mixed with 220 μl of assay buffer (100 mM Hepes-KOH, pH 7.8, 5 mM MgCl2, 10 mM KCl and 2 mM ATP). After 15 min of incubation at 30°C, the reaction was started by the addition of the fluorigenic substrate Suc-LLYY-NH-AMC (Calbiochem) and the release of amino-methyl-coumarin (360 nm ex/460 nm em) was monitored between 0 and 120 min, by Victor3 Multilabel Plate Readers (PerkinElmer). Protein concentration was measured using Protein Assay System (Bio-Rad) according to Bradford (Bradford, 1976), with serum albumin as standard.

### Nitrate reductase (NR) activity

The NR (EC 1.7.1.1) activity was measured according to previous work (Fortunato *et al*, 2019). Arabidopsis seedlings were ground in a mortar with 1:5 (w/v) extraction buffer (50 mM potassium phosphate pH 7,5; 1 mM EDTA, 1 mM DTT, 0,1 mM PMSF) and centrifuged at 20000 g for 15 min at 4 C. One volume of supernatants was incubated with five volumes of buffer (50 mM Hepes/KOH, pH 7.5, 0.04% (v/v) Triton X-100, 2 mM EDTA, 10 μM Na_2_MoO_4_, 20 μM flavin adenine dinucleotide, 0.5 mM DTT, 20 μM leupeptin, 20 mM potassium nitrate). The reaction was started by the addition of 0.6 mM NADH. After 15 min, 300 μL aliquots were kept, and the reaction was stopped by adding 25 μL of 0.6 mM zinc acetate. Finally, 300 μL 1% (w/v) sulfanilamide in 3 N HCl and 300 μL 0.02% (w/v) N(1-naphtyl)ethylendiamine dihydrochloride in 2.5% (v/v) H3PO4 were added. After 20 min, the absorbance was read at 540 nm.

### O_2_ consumption and AltOX activity analysis

Mitochondrial activity was determined through oxygen consumption and alternative oxidase (AltOX) activity analysis. Measurements were performed using a test version of the NextGen-O2k (Oroboros Instruments, Innsbruck) according to the methods developed previouysly (Vera-Vives *et al*, 2022). Mitochondria were isolated from 6-day-old seedlings grown on Murashige and Skoog medium supplemented with 2% (w/v) sucrose and 1.5% (w/v) Phyto-Agar, in the presence or absence of 550 µM Lin. Seedlings were homogenized in 300 mM sucrose, 60 mM MES, 10 mM EDTA, 25 mM Na4P2O7, 10 mM KH2PO4, 1 mM glycine, 50 mM sodium ascorbate, 20 mM cysteine, 1% Bovine Serum Albumin, 1% polyvinylpyrrolidone (pH 8) at a 1:6 (m/v) ratio. Homogenates were filtered (50 µm membrane; Miracloth) and centrifuged at 2500 *g* for 5 minutes at 4°C. The supernatant was centrifuged at 15000 g for 15 minutes at 4°C. The resulting mitochondria-enriched pellet was resuspended in 200 µL of 300 mM sucrose, 10 mM MES, 2 mM EDTA, 10 mM KH2PO4 (pH 7.5). Respiration rates were normalized to total protein content, which was measured using Infinite 200 PRO multimode plate reader (Tecan Group Ltd., Switzerland). The oxygen concentration was assessed in 2-ml measuring chambers at 22 °C with a 2-second frequency and samples were magnetically stirred at 750 rpm. Measurements were performed in the dark using 0.2 mg of total protein resuspended in Respiration Buffer (300 mM mannitol, 20 mM Hepes, 1 mM MgCl2, 20 mM KCl, 500 mM KH2PO4). Two measurements were done in parallel each time, taking advantage of the two chambers of the instrument. To stimulate respiration, malate (2 mM), pyruvate (5 mM), and thiamine pyrophosphate (TPP; 0.2 mM) were supplemented, followed by ADP (1 mM). n-propyl gallate (nPG; 0.5 mM) and Antimycin A (AA; 0.2 µM) were added to inhibit AltOX and Complex III activity, respectively. After each experiment the respiration chambers were washed with 100% ethanol and rinsed with deionized water at least six times. AltOX capacity was calculated as the percentage of AA-resistant respiration or nPG-inhibited respiration compared to total respiration, normalized to leak respiration. Maximal capacity was normalized to the mitochondrial marker voltage-dependent anion channel (VDAC). VDAC signals in Western Blot analysis of crude extracts were used to normalize each value of maximal respiration. Data were collected from a minimum of six replicates across at least four independent experiments.

### Nucleic acid analyses

For real-time Quantitative PCR (RT-qPCR) analyses, 1 µg of total RNA were processed using the iScript™ gDNA Clear cDNA Synthesis Kit (Bio-Rad; www.bio-rad.com), to achieve genomic DNA digestion and first-strand cDNA synthesis. RT-qPCR analyses were performed on a CFX96 Real-Time system (Bio-Rad; www.bio-rad.com), using primer pairs listed in **Table S7**. *PP2AA3* (*AT1G13320*) expression was probed as an internal reference (Czechowski *et al*, 2005). Raw data were analyzed with the Bio-Rad CFX Maestro 1.1 v 4.1 software (Bio-Rad; www.bio-rad.com). Samples were analyzed in triplicate.

### Chlorophyll-a fluorescence Measurements and Chlorophyll Quantification

The Chl-a fluorescence was measured in vivo employing the imaging Chl fluorometer (Walz Imaging PAM; walz.com), as previously described (Tadini *et al*, 2023). Dark-adapted seedlings allowed maximum quantum yield of PSII (*Fv*/*Fm*) measurements. The blue measuring light had the intensity set to 4 and saturating pulse intensity was also set to 4.

### Protoplast preparation and confocal microscopy

Protoplast isolation and transformation were performed as previously described (Yoo *et al*, 2007; Costa *et al*, 2012). Well-expanded rosette leaves from 18 DAS were cut into strips of 1–5 mm with a scalpel blade. Leaf tissue was incubated in an enzyme solution containing 1.25% (w/v) cellulase Onozuka R-10 (Duchefa) and 0.3% (w/v) Macerozyme R-10 (Duchefa) for 3 h at room temperature in the dark. The suspension was filtered through a 50 μm nylon mesh and washed with a solution containing 154 mM NaCl, 125 mM CaCl2, 5 mM KCl, 2 mM Mes (pH 5.7). For each protoplast transformation, 10 μg of plasmid DNA were employed. Protoplasts or leaves were observed through an up-right Nikon A1 confocal microscope. GFP channel excitation 485nm / detection 500-550nm. RFP channel excitation 560nm / detection 570-616nm. Chlorophyll channel excitation 485nm / detection 663-738nm.

## Author contribution

DDS, PP and LT conceptualized the study. GD, MM and CV produced proteomics data presented in this work. DDS, PM and AL performed data analysis and protein network reconstruction. AT, YLN and MZ performed respiration- and mitochondrial-related analyses. MDP and SF performed proteasome and nitrate reductase analyses. NJ, LTH and LT took care of mutant generation and performed the study of plastid-located hubs. Every author contributed to drafting the manuscript. PP, DDS, NJ and LT oversaw the study and finalized the manuscript. All authors approved its publication and accepted responsibility for the study.

## Acknowledgment

We are thankful to Antoni Mateu Vera Vives, Tomas Morosinotto and Alex Costa for the help and support in the experimental work, and to Nir Sade (University of Tel Aviv) for *Arabidopsis thaliana* amiR-CV seeds. We are also grateful to Norma Lattuada, Valerio Paravicini and Mario Beretta for technical assistance. Part of this work was carried out at NOLIMITS, an advanced imaging facility by the Università degli Studi di Milano. This study was carried out with the support of MUR—Ministero dell’Università e della Ricerca, grant number PRIN 2017-FBS8YN, entitled to PP.

## Highlights

Proteomics and systems biology approaches based on graph theory represent an effective combination of tools to identify proteins having key roles in plant cell adaptive responses

## Supplementary Figures

**Figure S1.**
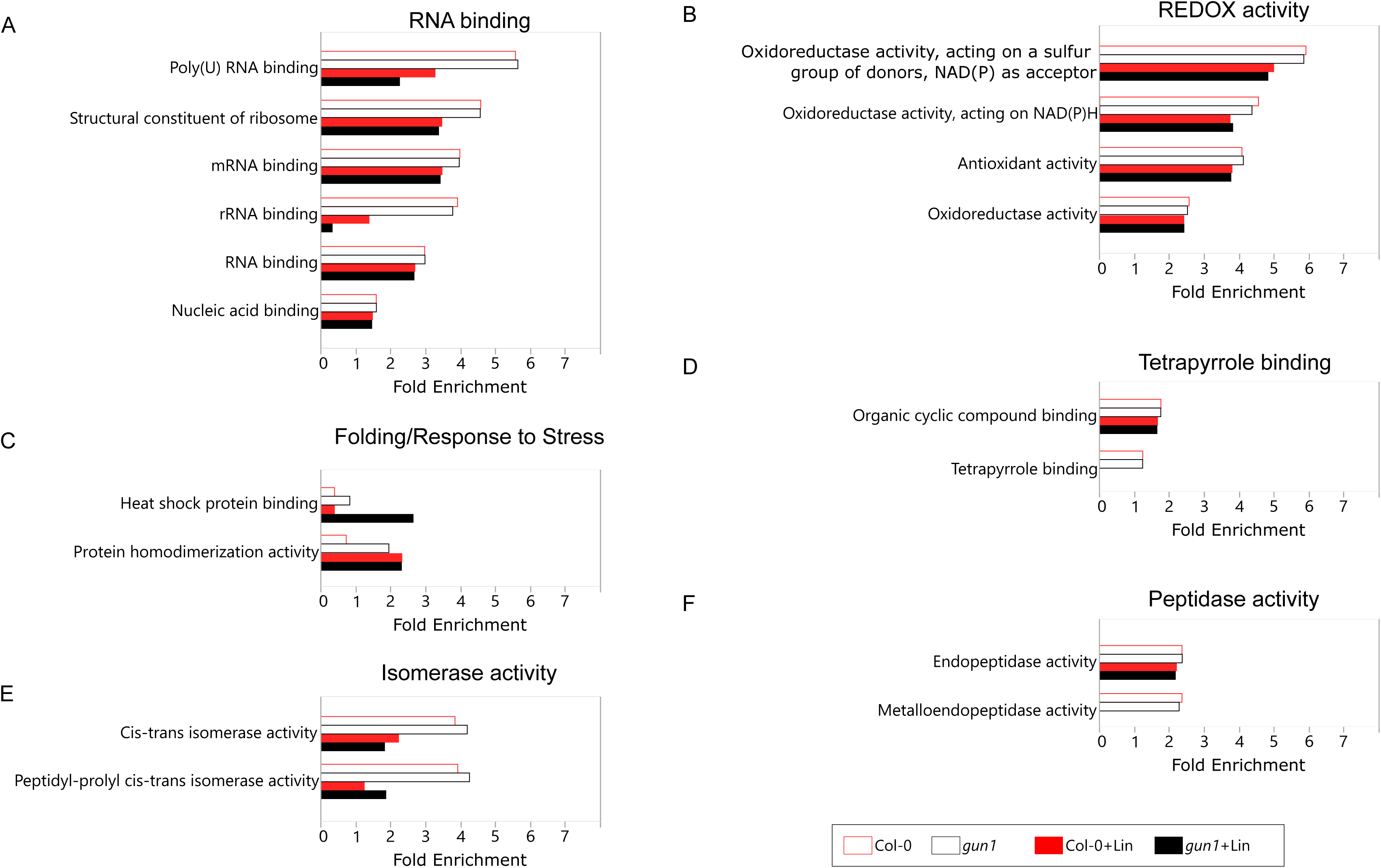
Molecular Functions (MFs) differentially enriched in *Arabidopsis thaliana* seedlings, Col-0 and *gun1* mutant, grown with or without lincomycin (+/-Lin) in the MS medium. MFs were retrieved from TAIR and PANTHER databases, while those differentially enriched were extracted by using LDA (n=9 per condition, p<0.01). **A**) RNA binding, **B**) REDOX activity, **C**) Folding/Response to Stress, **D**) Tetrapyrrole binding, **E**) Isomerase activity, **F**) Peptidase activity.

**Figure S2.**
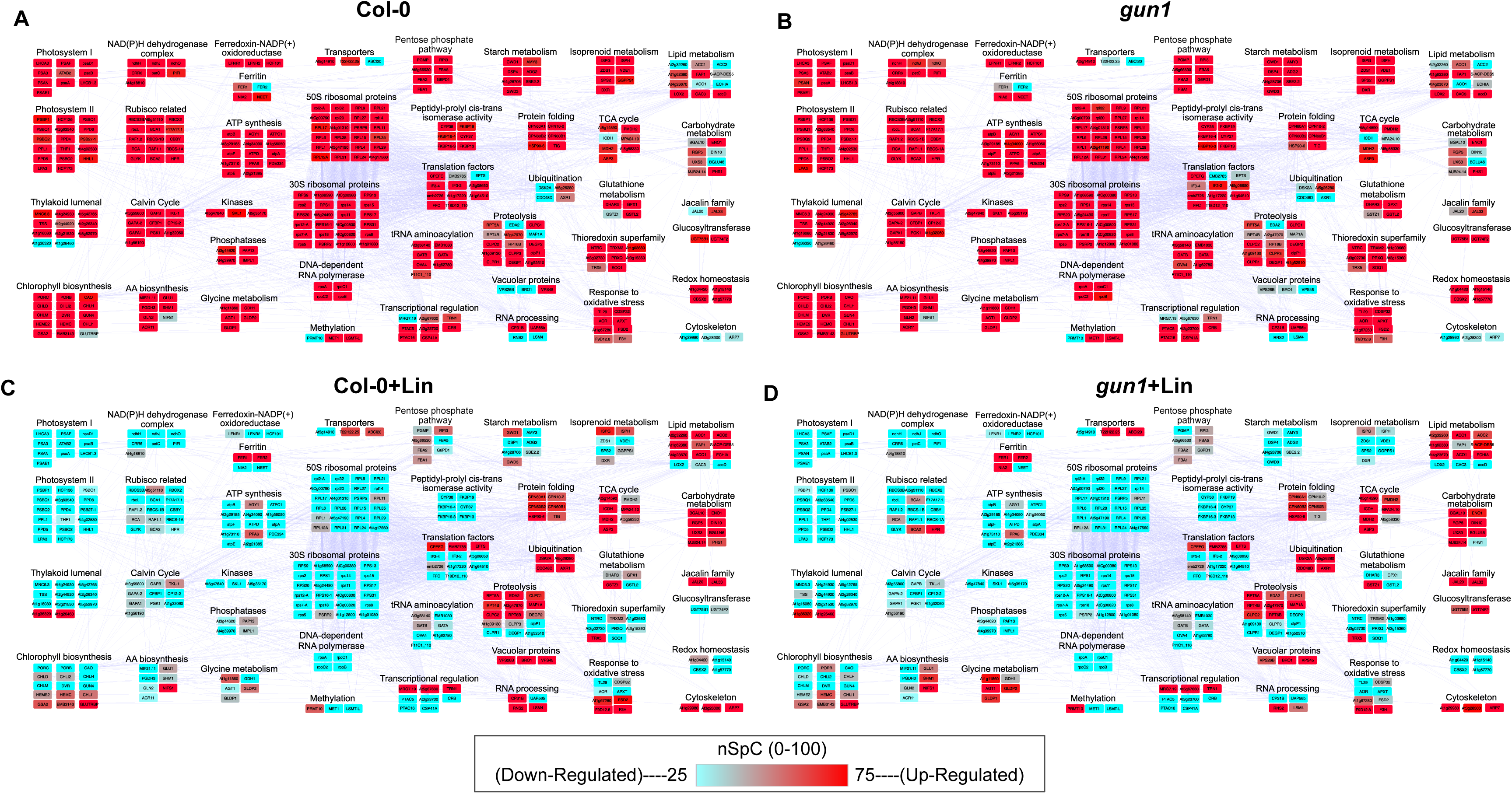
Protein-protein interaction (PPI) network modules affected by growth on Lin-containing medium and *gun1* mutation. The network was reconstructed from STRING db starting from high-confident differentially abundant proteins (DAPs) (p ≤ 0.001) The network counts 326 nodes and 2479 edges. Protein accumulation is represented by normalized spectral count (nSpC) values (range 0-100); specifically, blue and red nodes indicate down- and up-regulated proteins, respectively.

**Figure S3.**
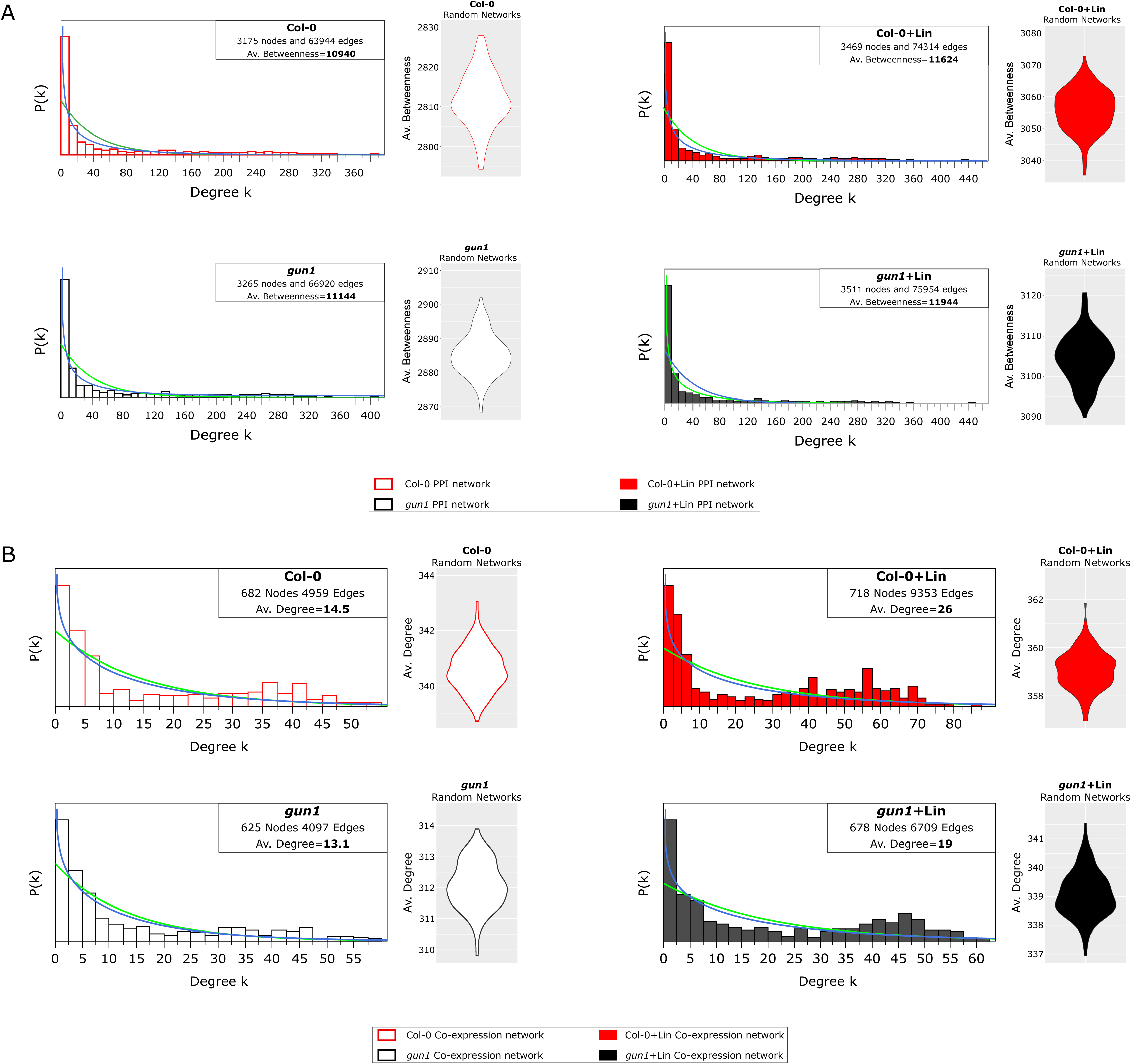
PPI and co-expression network models of *Arabidopsis thaliana* Col-0 and *gun1* seedlings, grown in absence and presence of lincomycin (-/+Lin). **A**) Degree distribution of Col-0, *gun1*, Col-0+Lin and *gun1*+Lin co-expression network models and violin plot reporting the average Betweenness in the corresponding random network models. **B**) Degree distribution of Col-0, *gun1*, Col-0+Lin and *gun1*+Lin co-expression network models and violin plot reporting the average Degree in the corresponding random network models.

**Figure S4.**
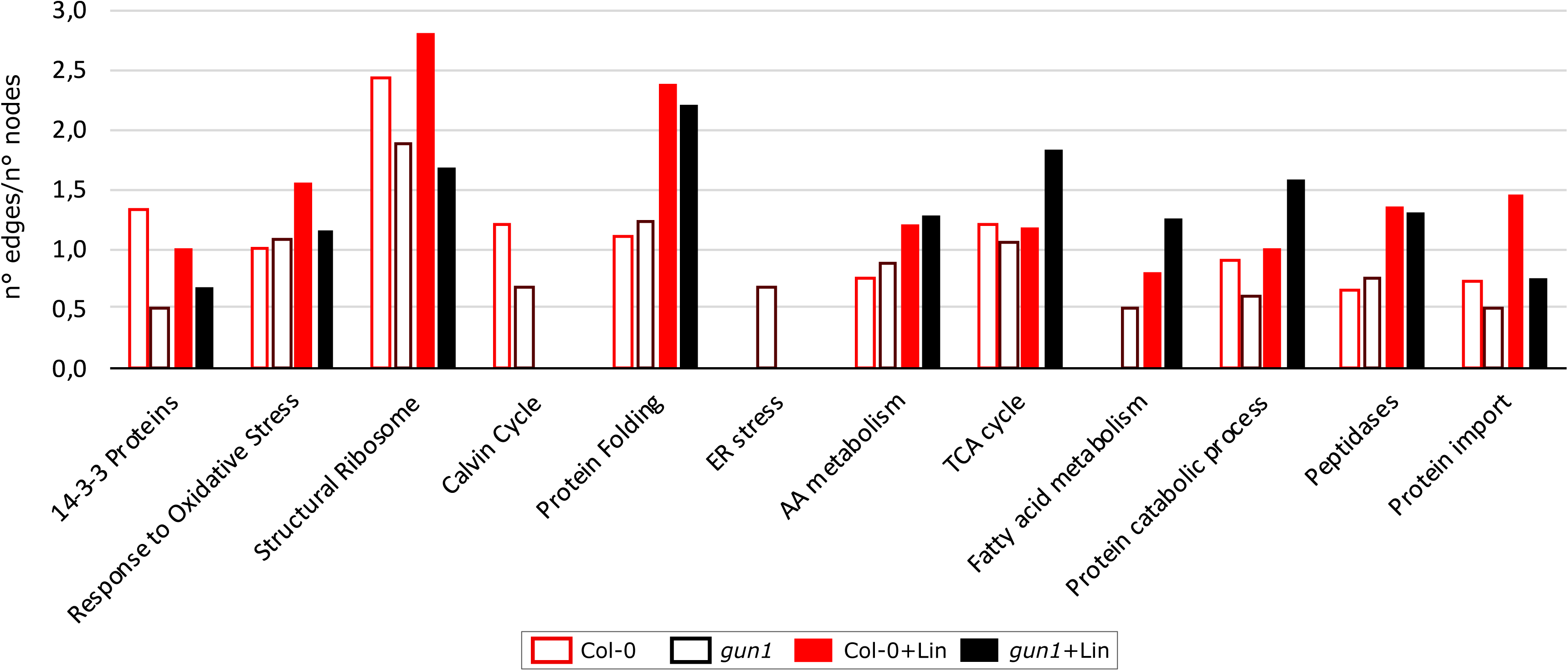
Biological Processes, Molecular Functions and Protein Complexes differentially correlated in Col-0 and *gun1* seedlings grown in absence or presence of Lin. For each of them, the degree of correlation was calculated by the ratio between the number of physical interactions with Spearman correlation > |0.75| and the number of proteins involved (n° edges/n°nodes), in Col-0, *gun1*, Col-0+Lin and *gun1*+Lin co-expression network models.

**Figure S5.**
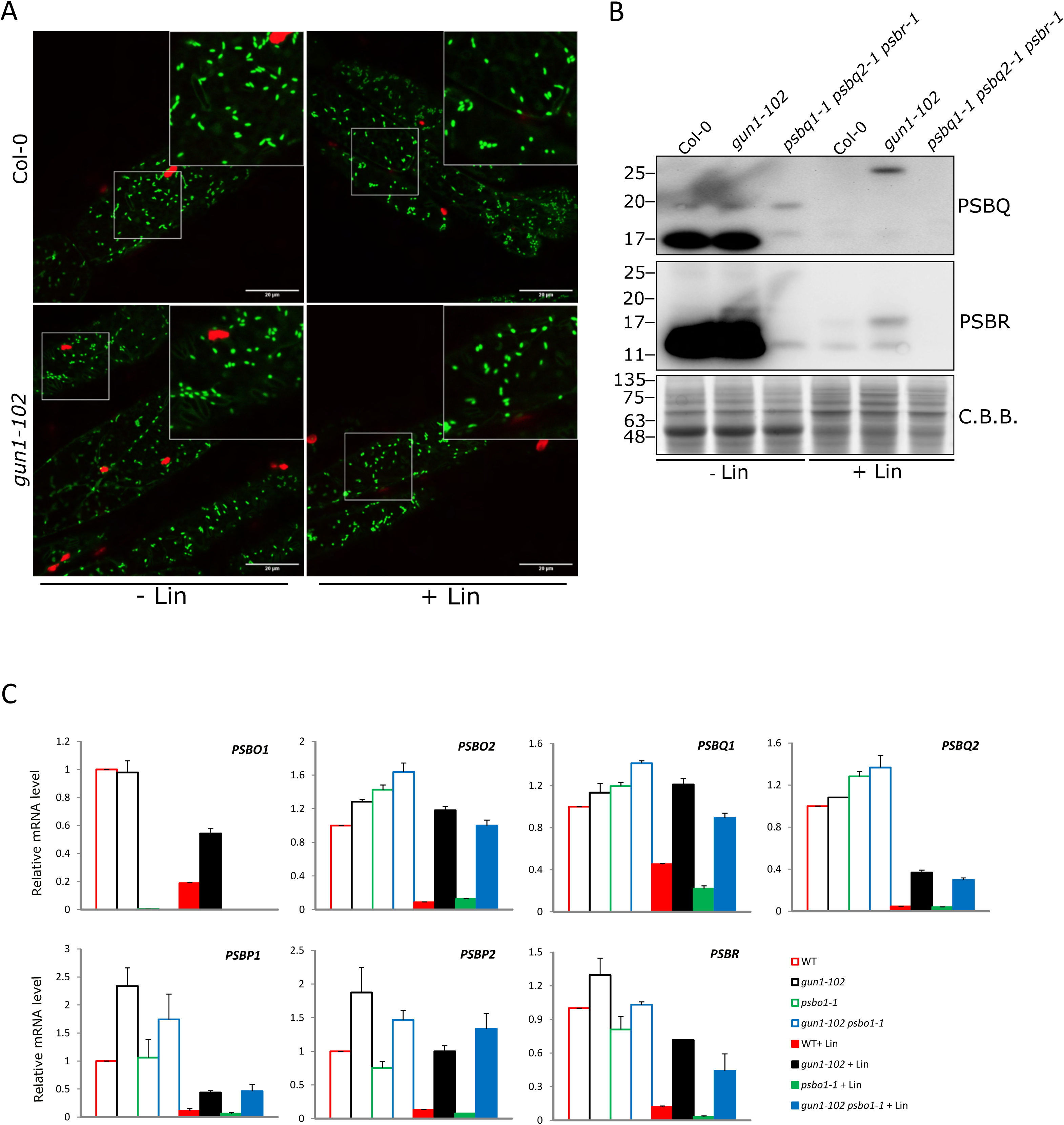
Mitochondrial morphology and OEC-gene expression analyses. **A**) Confocal microscopy observation of mitochondrial localized YFP protein (mTP-YFP) in Col-0 and *gun1-102* genetic backgrounds, grown in presence or absence of Lincomycin (-/+Lin; 550 µM). **B**) Immunoblot analyses of Col-0, *gun1-102* and *psbq1-1 psbq2-1 psbr-1* (- /+Lin 550 µM) total protein extract probed with PsbR- and PsbQ-specific antibodies. Coomassie Brilliant Blue (C.B.B.) staining of SDS-PAGE is shown as loading control. **C**) Real-time quantitative PCR on Oxygen Evolving Complex subunits *PsbO*, *PsbQ*, *PsbP* and *PsbR* in Col-0, *gun1-102*, *psbo1-1* and *gun1-102 psbo1-1* (-/+Lin 550 µM) genotypes.

**Figure S6.**
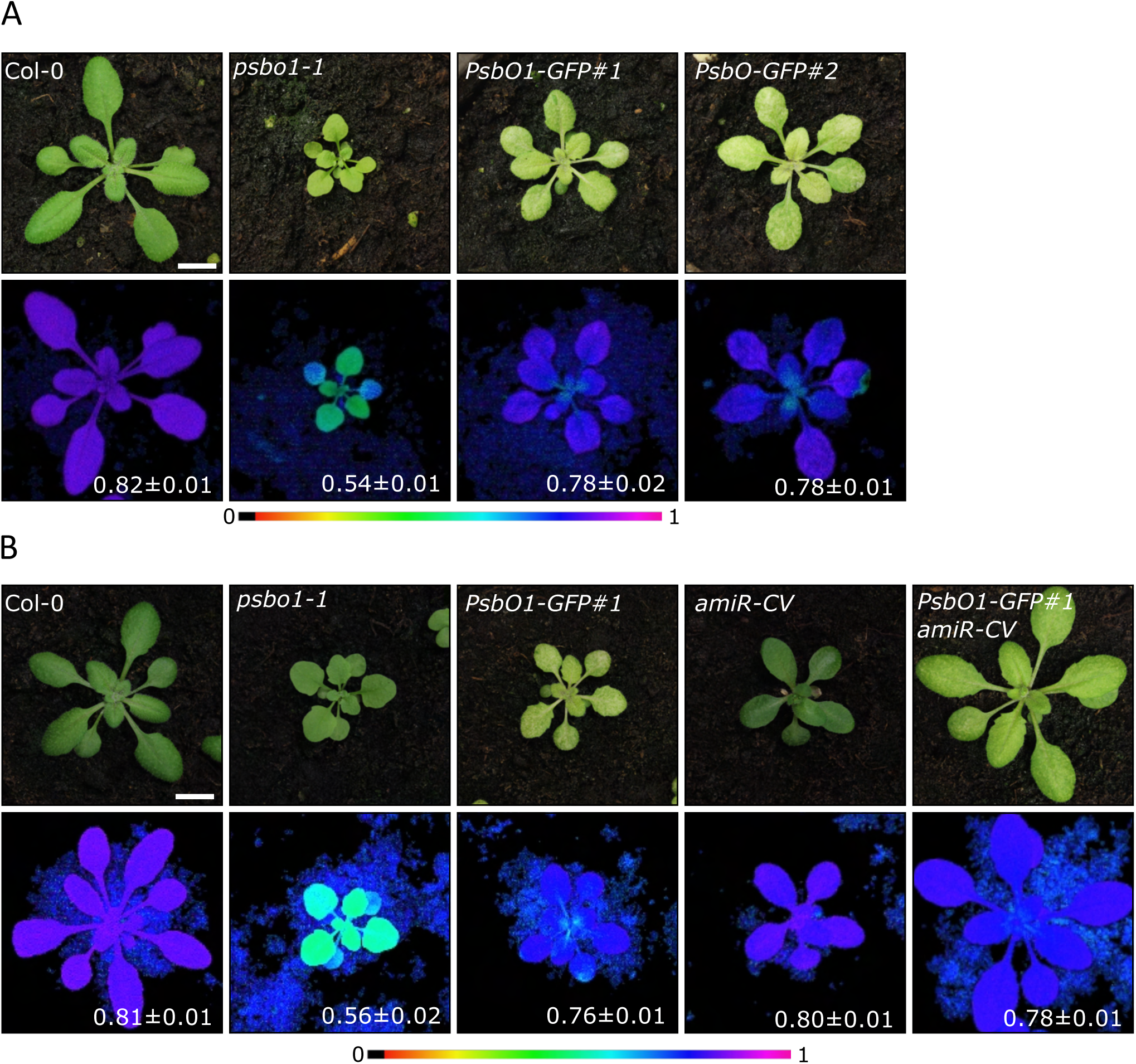
Visible phenotypes and photosynthetic performance of Arabidopsis Col-0, mutant and over-expressor lines. A) Immages of visible phenotypes displayed by 18 DAS plants of the indicated genotypes and Imaging PAM pictures representing in false colours the *Fv/Fm* parameter and the average ± SD values (n ≥ 4). The scale bar represents 1 cm. B) Pictures of the visible phenotypes displayed by 18 DAS plants of the indicated genotypes and Imaging PAM pictures representing in false colours the *Fv/Fm* parameter and the average ± SD values (n ≥ 4). The scale bar represents 1 cm.

**Figure S7.**
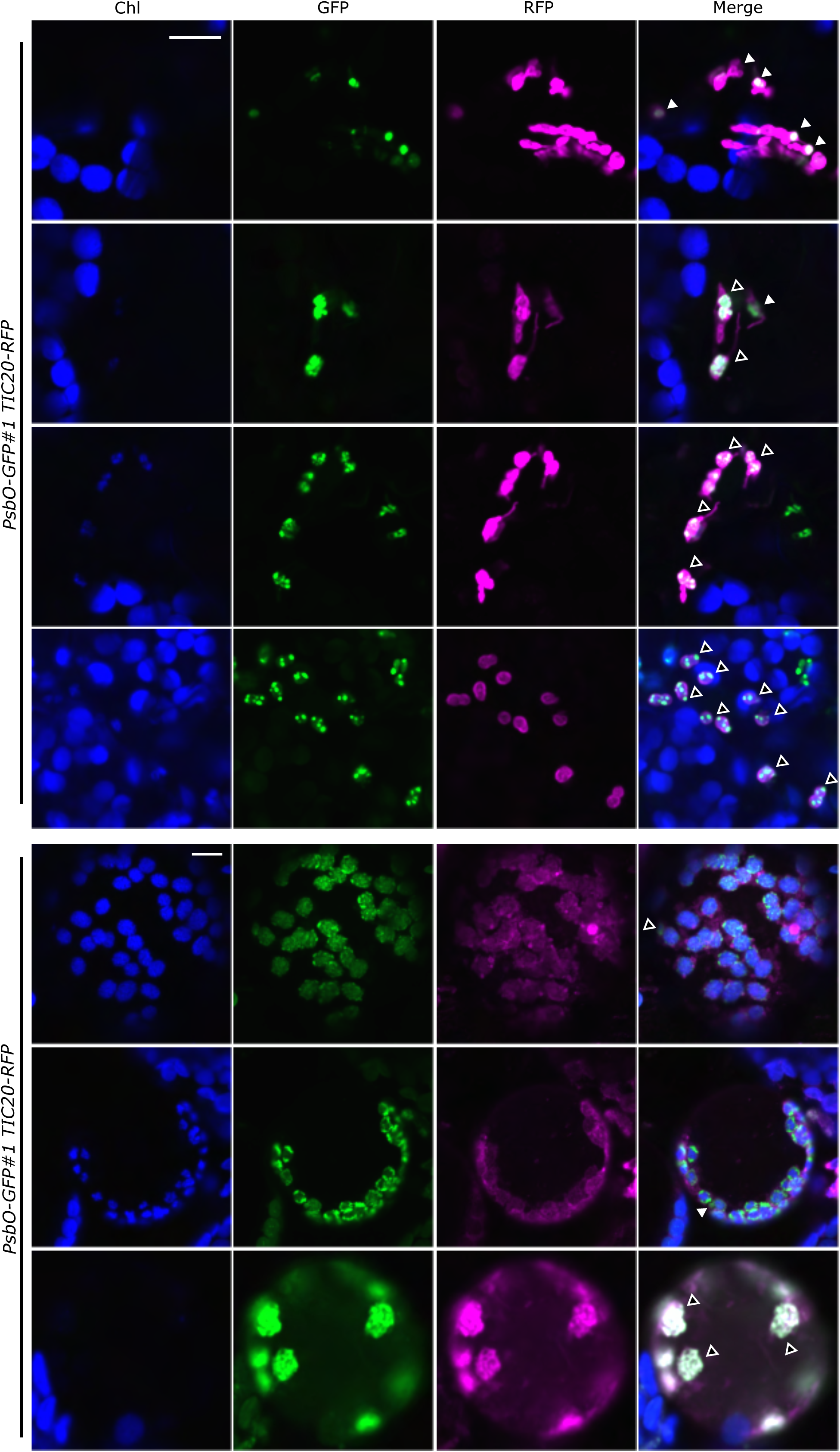
Fluorescence signals from chlorophylls (Chl, blue), GFP (green) and RFP (magenta) detected in mesophyll tissue (upper panel) or protoplasts (lower panel) obtained from *PsbO1-GFP#1 TIC20-RFP* line. Solid arrowheads indicate vesicles with GFP and RFP fluorescence, while empty arrowheads highlight chloroplasts showing GFP, RFP and Chl fluorescence. The scale bar represents 10 µm.

**Figure S8.**
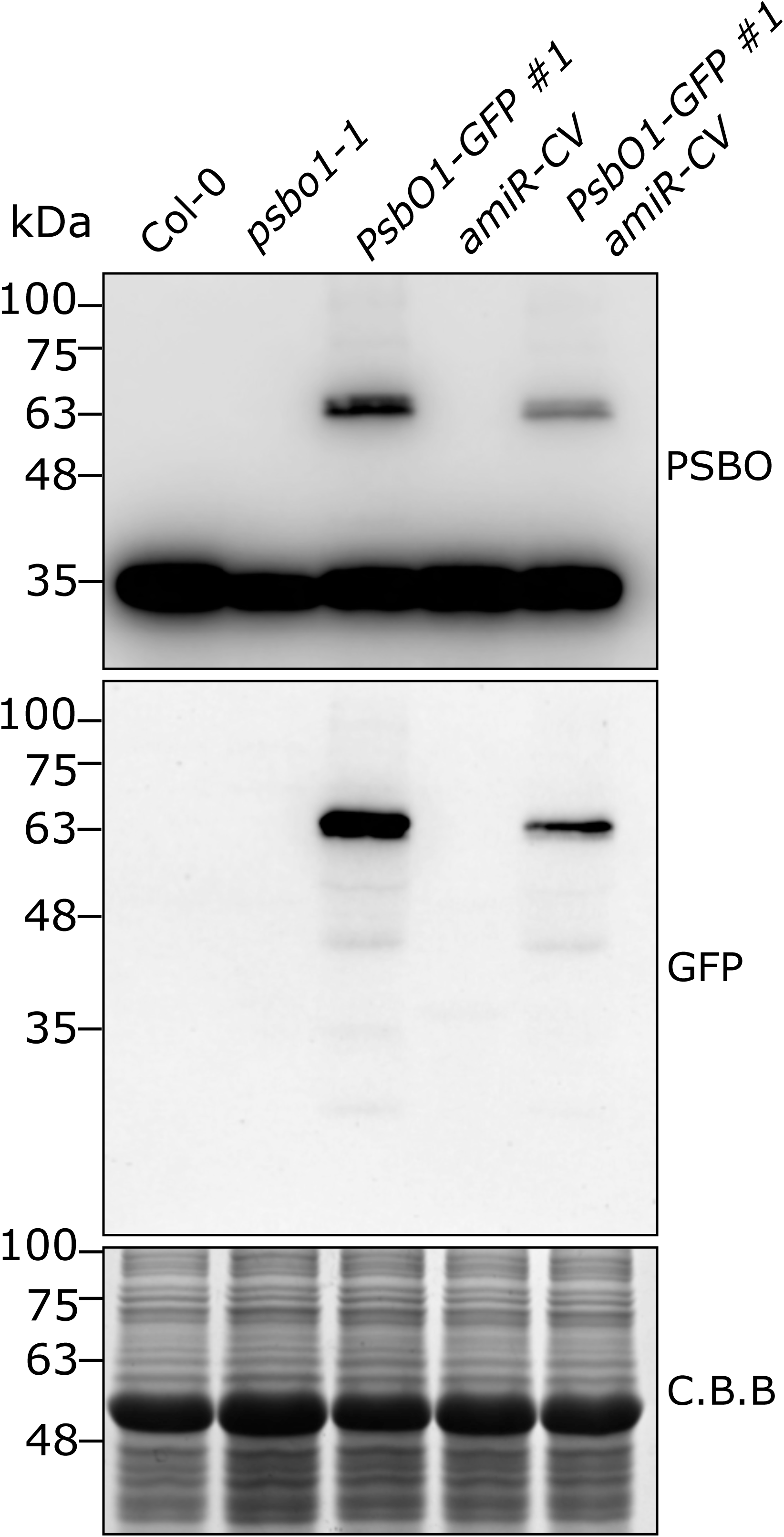
Immunoblot analyses on total protein extract from Col-0, *psbo1-1*, *PsbO1-GFP#1*, *amiR-CV* and *PsbO1-GFP#1 amiR-CV* double mutant grown on soil until 12 DAS probed with PsbO- and GFP-specific antibodies. Coomassie Brilliant Blue (C.B.B.) staining of SDS-PAGE is shown as loading control.

## Supplementary Tables

**Table S1.** Proteins identified in *Arabidopsis thaliana* Col-0 and *gun1* mutant, grown with and without lincomycin (+/- Lin). For each protein, the normalized Spectral Count (SpC) per biological replicate (rep) is shown, as well as the average SpC (Av.SpC) per condition. IF: Identification Frequency. In grey are highlighted common proteins found in all 36 replicates.

**Table S2**. GO Biological Processes (BPs) differentially enriched by comparing the enrichment profiles (by both Fold enrichment and n° proteins) obtained from Col-0, Col-0+Lin, *gun1* and *gun1*+Lin (n=9 per condition). BPs were retrieved from TAIR and PANTHER databases, while those differentially enriched were extracted by Linear Discriminant Analysis (LDA, p≤0.01, F rat =LN(SpC1/SpC2) is shown.

**Table S3**. A) Differentially Abundant Proteins (DAPs) in Arabidopsis thaliana Col-0 and *gun1* mutant, grown with and without lincomycin (+/-Lin). Proteins were selected by LDA (P<0.05). Most significant DAPs (P <0.01) are highlighted in bold. A fold change calculated as FC=LN(SpC1/SpC2) is shown; specifically, positive values (red) indicate proteins up-regulated in the first condition (red), while negative values (blue) indicate proteins up-regulated in the second condition (blue). B) Subset of proteins listed in A in which the absolute protein abundance has been estimated by normalization using the average amino acid molecular weight.

**Table S4**. PPI network hubs characterizing the proteome obtained from *Arabidopsis thaliana* Col-0 and *gun1* seedlings grown on MS medium with and without lincomycin (+/-Lin). Proteins were defined hubs if Betweenness, Centroid and Bridging values were higher than the average values calculated on whole network models (in bold, black); while in red centrality values not exceeding the imposed thresholds are indicated. In yellow are highlighted centralities values used to identify hubs characterizing specific phenotypes. For each hub, the average Spectral Count and the fold change (if DAP, too) are shown. A) Hubs in all conditions. B) Hubs in Col-0. C) Hubs in *gun1*. D) Hubs in Col-0+Lin. E) Hubs in *gun1*+Lin. F) Hubs characterizing Lin growth. G) Hubs characterizing *gun1*.

**Table S5**. Co-expression network hubs characterizing the proteome obtained from *Arabidopsis thaliana* Col-0 and *gun1* seedlings grown on MS medium with and without lincomycin (+/-Lin). Proteins were defined hubs if the Degree value was higher than the average values calculated on whole network models (in bold, black) and it was less than the average in the other conditions. For each protein, GO Cellular Component is shown. In yellow are highlighted Degree values used to identify differentially co-expressed hubs characterizing specific phenotypes. A) Hubs in Col-0, B) Hubs in *gun1*, A) Hubs in Col-0+Lin, D) Hubs in *gun1*+Lin. E) Hubs in Col-0 and Col-0+Lin. F) Hubs in *gun1* and *gun1*+Lin, G) Hubs Col-0 and *gun1*, H) Hubs in Col-0+Lin and *gun1*+Lin.

**Table S6**. Tryptic peptides identified by LC-MS/MS analysis in higher molecular weight PsbO bands (+ and * bands, respectively) identified by immunoblot analyses on total protein extract from seedlings grown in presence of 550 µM Lincomycin probed with PsbO antibody. In blue are peptides mapping on chloroplast transit peptide sequence (residues 1 to 58), in red are peptides mapping on thylakoid transit peptides sequence (residues 59 to 85).

**Table S7**. Oligonucleotide sequences employed for RT-qPCR and cloning.

## Bibliography

Van Aken O & Van Breusegem F (2015) Licensed to Kill: Mitochondria, Chloroplasts, and Cell Death. Trends Plant Sci 20: 754–766

Van Aken O & Pogson BJ (2017) Convergence of mitochondrial and chloroplastic ANAC017/PAP-dependent retrograde signalling pathways and suppression of programmed cell death. Cell Death Differ 24: 955–960

Van Aken O, Zhang B, Law S, Narsai R & Whelan J (2013) AtWRKY40 and AtWRKY63 Modulate the Expression of Stress-Responsive Nuclear Genes Encoding Mitochondrial and Chloroplast Proteins. Plant Physiol 162: 254– 271

Allahverdiyeva Y, Suorsa M, Rossi F, Pavesi A, Kater MM, Antonacci A, Tadini L, Pribil M, Schneider A, Wanner G, et al (2013) Arabidopsis plants lacking PsbQ and PsbR subunits of the oxygen-evolving complex show altered PSII super-complex organization and short-term adaptive mechanisms. The Plant Journal 75: 671–684

Anjum NA, Sharma P, Gill SS, Hasanuzzaman M, Khan EA, Kachhap K, Mohamed AA, Thangavel P, Devi GD, Vasudhevan P, et al (2016) Catalase and ascorbate peroxidase—representative H2O2-detoxifying heme enzymes in plants. Environmental Science and Pollution Research 23: 19002–19029

Assenov Y, Ramírez F, Schelhorn S-E, Lengauer T & Albrecht M (2008) Computing topological parameters of biological networks. Bioinformatics 24: 282–284

Berardini TZ, Reiser L, Li D, Mezheritsky Y, Muller R, Strait E & Huala E (2015) The arabidopsis information resource: Making and mining the “gold standard” annotated reference plant genome. genesis 53: 474–485

Bjornson M, Balcke GU, Xiao Y, de Souza A, Wang J, Zhabinskaya D, Tagkopoulos I, Tissier A & Dehesh K (2017) Integrated omics analyses of retrograde signaling mutant delineate interrelated stress-response strata. The Plant Journal 91: 70–84

Blanco NE, Guinea-Díaz M, Whelan J & Strand Å (2014) Interaction between plastid and mitochondrial retrograde signalling pathways during changes to plastid redox status. Philosophical Transactions of the Royal Society B: Biological Sciences 369: 20130231

Boex-Fontvieille E, Rustgi S, Reinbothe S & Reinbothe C (2015) A Kunitz-type protease inhibitor regulates programmed cell death during flower development in *Arabidopsis thaliana*. J Exp Bot 66: 6119–6135

Bradford M (1976) A Rapid and Sensitive Method for the Quantitation of Microgram Quantities of Protein Utilizing the Principle of Protein-Dye Binding. Anal Biochem 72: 248–254

Chotewutmontri P & Barkan A (2018) Multilevel effects of light on ribosome dynamics in chloroplasts program genome-wide and psbA-specific changes in translation. PLoS Genet 14: e1007555

De Clercq I, Vermeirssen V, Van Aken O, Vandepoele K, Murcha MW, Law SR, Inzé A, Ng S, Ivanova A, Rombaut D, et al (2013) The Membrane-Bound NAC Transcription Factor ANAC013 Functions in Mitochondrial Retrograde Regulation of the Oxidative Stress Response in *Arabidopsis*. Plant Cell 25: 3472–3490

Colombo M, Tadini L, Peracchio C, Ferrari R & Pesaresi P (2016) GUN1, a Jack-Of-All-Trades in Chloroplast Protein Homeostasis and Signaling. Front Plant Sci 7

Costa A, Gutla PVK, Boccaccio A, Scholz-Starke J, Festa M, Basso B, Zanardi I, Pusch M, Schiavo F Lo, Gambale F, et al (2012) The *Arabidopsis* central vacuole as an expression system for intracellular transporters: functional characterization of the Cl ^−^ /H ^+^ exchanger CLC-7. J Physiol 590: 3421–3430

Czechowski T, Stitt M, Altmann T, Udvardi MK & Scheible W-R (2005) Genome-Wide Identification and Testing of Superior Reference Genes for Transcript Normalization in Arabidopsis. Plant Physiol 139: 5–17

Damiano F, Testini M, Tocci R, Gnoni G V. & Siculella L (2018) Translational control of human acetyl-CoA carboxylase 1 mRNA is mediated by an internal ribosome entry site in response to ER stress, serum deprivation or hypoxia mimetic CoCl2. Biochimica et Biophysica Acta (BBA) - Molecular and Cell Biology of Lipids 1863: 388–398

Domingo G, Chiodaroli L, Parola S, Marsoni M, Bracale M & Vannini C (2023) Proteomic characterization of Shiitake (Lentinula edodes) post-harvest fruit bodies grown on hardwood logs and isolation of an antibacterial serine protease inhibitor. Fungal Biol 127: 881–890

Doncheva NT, Morris JH, Gorodkin J & Jensen LJ (2019) Cytoscape StringApp: Network Analysis and Visualization of Proteomics Data. J Proteome Res 18: 623–632

Estavillo GM, Crisp PA, Pornsiriwong W, Wirtz M, Collinge D, Carrie C, Giraud E, Whelan J, David P, Javot H, et al (2011) Evidence for a SAL1-PAP Chloroplast Retrograde Pathway That Functions in Drought and High Light Signaling in *Arabidopsis*. Plant Cell 23: 3992–4012

Fang X, Zhao G, Zhang S, Li Y, Gu H, Li Y, Zhao Q & Qi Y (2019) Chloroplast-to-Nucleus Signaling Regulates MicroRNA Biogenesis in Arabidopsis. Dev Cell 48: 371–382.e4

Fortunato, Nigro, Paradiso, Cucci, Lacolla, Trani, Agrimi, Blanco, de Pinto & Gadaleta (2019) Nitrogen Metabolism at Tillering Stage Differently Affects the Grain Yield and Grain Protein Content in Two Durum Wheat Cultivars. Diversity (Basel) 11: 186

Fortunato S, Lasorella C, Tadini L, Jeran N, Vita F, Pesaresi P & de Pinto MC (2022) GUN1 involvement in the redox changes occurring during biogenic retrograde signaling. Plant Science 320: 111265

Fryer MJ, Ball L, Oxborough K, Karpinski S, Mullineaux PM & Baker NR (2003) Control of *Ascorbate Peroxidase 2* expression by hydrogen peroxide and leaf water status during excess light stress reveals a functional organisation of *Arabidopsis* leaves. The Plant Journal 33: 691–705

Giraud E, Van Aken O, Ho LHM & Whelan J (2009) The Transcription Factor ABI4 Is a Regulator of Mitochondrial Retrograde Expression of *ALTERNATIVE OXIDASE1a*. Plant Physiol 150: 1286–1296

Hatsugai N, Kuroyanagi M, Yamada K, Meshi T, Tsuda S, Kondo M, Nishimura M & Hara-Nishimura I (2004) A Plant Vacuolar Protease, VPE, Mediates Virus-Induced Hypersensitive Cell Death. Science (1979) 305: 855–858

Havé M, Balliau T, Cottyn-Boitte B, Dérond E, Cueff G, Soulay F, Lornac A, Reichman P, Dissmeyer N, Avice J-C, et al (2018) Increases in activity of proteasome and papain-like cysteine protease in Arabidopsis autophagy mutants: back-up compensatory effect or cell-death promoting effect? J Exp Bot 69: 1369–1385

Hernández-Verdeja T, Vuorijoki L, Jin X, Vergara A, Dubreuil C & Strand Å (2022) GENOMES UNCOUPLED1 plays a key role during the de-etiolation process in Arabidopsis. New Phytologist 235: 188–203

Huang J, Wang M-M, Jiang Y, Bao Y-M, Huang X, Sun H, Xu D-Q, Lan H-X & Zhang H-S (2008) Expression analysis of rice A20/AN1-type zinc finger genes and characterization of ZFP177 that contributes to temperature stress tolerance. Gene 420: 135–144

Huang X, von Rad U & Durner J (2002) Nitric oxide induces transcriptional activation of the nitric oxide-tolerant alternative oxidase in Arabidopsis suspension cells. Planta 215: 914–923

Jan M, Liu Z, Rochaix J-D & Sun X (2022) Retrograde and anterograde signaling in the crosstalk between chloroplast and nucleus. Front Plant Sci 13

Jeong H, Mason SP, Barabási A-L & Oltvai ZN (2001) Lethality and centrality in protein networks. Nature 411: 41– 42

Jeran N, Rotasperti L, Frabetti G, Calabritto A, Pesaresi P & Tadini L (2021) The PUB4 E3 ubiquitin ligase is responsible for the variegated phenotype observed upon alteration of chloroplast protein homeostasis in arabidopsis cotyledons. Genes (Basel) 12: 1387

Jia Y, Tian H, Zhang S, Ding Z & Ma C (2019) GUN1-Interacting Proteins Open the Door for Retrograde Signaling. Trends Plant Sci 24: 884–887

Jiang T, Oh ES, Bonea D & Zhao R (2017) HSP90C interacts with PsbO1 and facilitates its thylakoid distribution from chloroplast stroma in Arabidopsis. PLoS One 12: e0190168

Kemp C, Coleman A, Wells G & Parry G (2015) Overexpressing components of the nuclear transport apparatus causes severe growth symptoms in tobacco leaves. Plant Signal Behav 10: e1000103

Khator K & Shekhawat GS (2020) Cd- and Cu-induced phytotoxicity on 2–3 leaf stage of Cyamopsis tetragonoloba and its regulation by nitrate reductase and ROS quenching enzyme. Acta Physiol Plant 42: 120

Klasek L, Inoue K & Theg SM (2020) Chloroplast Chaperonin-Mediated Targeting of a Thylakoid Membrane Protein. Plant Cell 32: 3884–3901

Koh E, Carmieli R, Mor A & Fluhr R (2016) Singlet Oxygen-Induced Membrane Disruption and Serpin-Protease Balance in Vacuolar-Driven Cell Death. Plant Physiol 171: 1616–1625

Köhler D, Helm S, Agne B & Baginsky S (2016) Importance of Translocon Subunit Tic56 for rRNA Processing and Chloroplast Ribosome Assembly. Plant Physiol 172: 2429–2444

Koussevitzky S, Nott A, Mockler TC, Hong F, Sachetto-martins G, Surpin M, Lim J, Mittler R & Chory J (2007) Signals from Chloroplasts Converge to Regulate Nuclear Gene Expression. Science (1979) 316: 715–719

Lasorella C, Fortunato S, Dipierro N, Jeran N, Tadini L, Vita F, Pesaresi P & de Pinto MC (2022) Chloroplast-localized GUN1 contributes to the acquisition of basal thermotolerance in Arabidopsis thaliana. Front Plant Sci 13

Lemke MD, Fisher KE, Kozlowska MA, Tano DW & Woodson JD (2021) The core autophagy machinery is not required for chloroplast singlet oxygen-mediated cell death in the Arabidopsis thaliana plastid ferrochelatase two mutant. BMC Plant Biol 21: 1–20

Lemke MD & Woodson JD (2022) Targeted for destruction: degradation of singlet oxygen-damaged chloroplasts. Plant Signal Behav 17

Llamas E, Pulido P & Rodriguez-Concepcion M (2017) Interference with plastome gene expression and Clp protease activity in Arabidopsis triggers a chloroplast unfolded protein response to restore protein homeostasis. PLoS Genet 13: 1–28

Lundin B, Hansson M, Schoefs B, Vener A V. & Spetea C (2007) The Arabidopsis PsbO2 protein regulates dephosphorylation and turnover of the photosystem II reaction centre D1 protein. The Plant Journal 49: 528–539

Maere S, Heymans K & Kuiper M (2005) BiNGO: a Cytoscape plugin to assess overrepresentation of Gene Ontology categories in Biological Networks. Bioinformatics 21: 3448–3449

Manbir, Singh P, Kumari A & Gupta KJ (2022) Alternative oxidase plays a role in minimizing ROS and RNS produced under salinity stress in *Arabidopsis thaliana*. Physiol Plant 174

Marino G, Naranjo B, Wang J, Penzler JF, Kleine T & Leister D (2019) Relationship of GUN1 to FUG1 in chloroplast protein homeostasis. Plant Journal 99: 521–535

Maruta T, Noshi M, Tanouchi A, Tamoi M, Yabuta Y, Yoshimura K, Ishikawa T & Shigeoka S (2012) H2O2-triggered Retrograde Signaling from Chloroplasts to Nucleus Plays Specific Role in Response to Stress. Journal of Biological Chemistry 287: 11717–11729

Mi H, Ebert D, Muruganujan A, Mills C, Albou LP, Mushayamaha T & Thomas PD (2021) PANTHER version 16: A revised family classification, tree-based classification tool, enhancer regions and extensive API. Nucleic Acids Res 49: D394–D403

Milanesi R, Coccetti P & Tripodi F (2020) The Regulatory Role of Key Metabolites in the Control of Cell Signaling. Biomolecules 10: 862

Millar AH & Day DA (1996) Nitric oxide inhibits the cytochrome oxidase but not the alternative oxidase of plant mitochondria. FEBS Lett 398: 155–158

Mitschke J, Fuss J, Blum T, Höglund A, Reski R, Kohlbacher O & Rensing SA (2009) Prediction of dual protein targeting to plant organelles. New Phytologist 183: 224–236

Moreno JC, Mi J, Alagoz Y & Al-Babili S (2021) Plant apocarotenoids: from retrograde signaling to interspecific communication. The Plant Journal 105: 351–375

Muntz K (2007) Protein dynamics and proteolysis in plant vacuoles. J Exp Bot 58: 2391–2407

Ng S, De Clercq I, Van Aken O, Law SR, Ivanova A, Willems P, Giraud E, Van Breusegem F & Whelan J (2014) Anterograde and Retrograde Regulation of Nuclear Genes Encoding Mitochondrial Proteins during Growth, Development, and Stress. Mol Plant 7: 1075–1093

Nishizawa A, Yabuta Y, Yoshida E, Maruta T, Yoshimura K & Shigeoka S (2006) Arabidopsis heat shock transcription factor A2 as a key regulator in response to several types of environmental stress. Plant Journal 48: 535–547

Ono M, Isono K, Sakata Y & Taji T (2021a) CATALASE2 plays a crucial role in long-term heat tolerance of Arabidopsis thaliana. Biochem Biophys Res Commun 534: 747–751

Ono S, Suzuki S, Ito D, Tagawa S, Shiina T & Masuda S (2021b) Plastidial (p)ppGpp Synthesis by the Ca2+-Dependent RelA–SpoT Homolog Regulates the Adaptation of Chloroplast Gene Expression to Darkness in Arabidopsis. Plant Cell Physiol 61: 2077–2086

Pan Q-N, Geng C-C, Li D-D, Xu S-W, Mao D-D, Umbreen S, Loake GJ & Cui B-M (2019) Nitrate Reductase-Mediated Nitric Oxide Regulates the Leaf Shape in Arabidopsis by Mediating the Homeostasis of Reactive Oxygen Species. Int J Mol Sci 20: 2235

Paradiso A, Domingo G, Blanco E, Buscaglia A, Fortunato S, Marsoni M, Scarcia P, Caretto S, Vannini C & de Pinto MC (2020) Cyclic AMP mediates heat stress response by the control of redox homeostasis and ubiquitin-proteasome system. Plant Cell Environ 43: 2727–2742

Parcerisa IL, Rosano GL & Ceccarelli EA (2020) Biochemical characterization of ClpB3, a chloroplastic disaggregase from Arabidopsis thaliana. Plant Mol Biol 104: 451–465

Park S & Rodermel SR (2004) Mutations in ClpC2/Hsp100 suppress the requirement for FtsH in thylakoid membrane biogenesis. Proceedings of the National Academy of Sciences 101: 12765–12770

Parker N, Wang Y & Meinke D (2014) Natural Variation in Sensitivity to a Loss of Chloroplast Translation in Arabidopsis. Plant Physiol 166: 2013–2027

Parker N, Wang Y & Meinke D (2016) Analysis of Arabidopsis Accessions Hypersensitive to a Loss of Chloroplast Translation. Plant Physiol 172: 1862–1875

Perez de Souza L, Garbowicz K, Brotman Y, Tohge T & Fernie AR (2020) The Acetate Pathway Supports Flavonoid and Lipid Biosynthesis in Arabidopsis. Plant Physiol 182: 857–869

Perez-Riverol Y, Bai J, Bandla C, García-Seisdedos D, Hewapathirana S, Kamatchinathan S, Kundu DJ, Prakash A, Frericks-Zipper A, Eisenacher M, et al (2022) The PRIDE database resources in 2022: a hub for mass spectrometry-based proteomics evidences. Nucleic Acids Res 50: D543–D552

Pesaresi P, Masiero S, Eubel H, Braun H-P, Bhushan S, Glaser E, Salamini F & Leister D (2006) Nuclear Photosynthetic Gene Expression Is Synergistically Modulated by Rates of Protein Synthesis in Chloroplasts and Mitochondria. Plant Cell 18: 970–991

Richter AS, Nägele T, Grimm B, Kaufmann K, Schroda M, Leister D & Kleine T (2023) Retrograde signaling in plants: A critical review focusing on the GUN pathway and beyond. Plant Commun 4: 100511

Salgado I, Carmen Martínez M, Oliveira HC & Frungillo L (2013) Nitric oxide signaling and homeostasis in plants: a focus on nitrate reductase and S-nitrosoglutathione reductase in stress-related responses. Brazilian Journal of Botany 36: 89–98

Sanmartín M, Ordóñez A, Sohn EJ, Robert S, Sánchez-Serrano JJ, Surpin MA, Raikhel N V. & Rojo E (2007) Divergent functions of VTI12 and VTI11 in trafficking to storage and lytic vacuoles in *Arabidopsis*. Proceedings of the National Academy of Sciences 104: 3645–3650

Scardoni G, Tosadori G, Faizan M, Spoto F, Fabbri F & Laudanna C (2015) Biological network analysis with CentiScaPe: centralities and experimental dataset integration. F1000Res 3: 139

Schägger H & von Jagow G (1987) Tricine-sodium dodecyl sulfate-polyacrylamide gel electrophoresis for the separation of proteins in the range from 1 to 100 kDa. Anal Biochem 166: 368–379

Shahriari M, Keshavaiah C, Scheuring D, Sabovljevic A, Pimpl P, Häusler RE, Hülskamp M & Schellmann S (2010) The AAA-type ATPase AtSKD1 contributes to vacuolar maintenance of Arabidopsis thaliana. The Plant Journal 64: 71–85

Shapiguzov A, Vainonen JP, Hunter K, Tossavainen H, Tiwari A, Järvi S, Hellman M, Aarabi F, Alseekh S, Wybouw B, et al (2019) Arabidopsis RCD1 coordinates chloroplast and mitochondrial functions through interaction with ANAC transcription factors. Elife 8

Shimada T, Takagi J, Ichino T, Shirakawa M & Hara-Nishimura I (2018) Plant Vacuoles. Annu Rev Plant Biol 69: 123–145

Di Silvestre D, Bergamaschi A, Bellini E & Mauri P (2018) Large Scale Proteomic Data and Network-Based Systems Biology Approaches to Explore the Plant World. Proteomes 6: 27

Di Silvestre D, Passignani G, Rossi R, Ciuffo M, Turina M, Vigani G & Mauri PL (2022) Presence of a Mitovirus Is Associated with Alteration of the Mitochondrial Proteome, as Revealed by Protein–Protein Interaction (PPI) and Co-Expression Network Models in Chenopodium quinoa Plants. Biology (Basel) 11: 95

Di Silvestre D, Vigani G, Mauri P, Hammadi S, Morandini P & Murgia I (2021) Network Topological Analysis for the Identification of Novel Hubs in Plant Nutrition. Front Plant Sci 12

Srivastava R, Deng Y, Shah S, Rao AG & Howell SH (2013) BINDING PROTEIN Is a Master Regulator of the Endoplasmic Reticulum Stress Sensor/Transducer bZIP28 in *Arabidopsis*. Plant Cell 25: 1416–1429

Su G, Morris JH, Demchak B & Bader GD (2014) Biological Network Exploration with Cytoscape 3. Curr Protoc Bioinformatics 47

Suorsa M & Aro E-M (2007) Expression, assembly and auxiliary functions of photosystem II oxygen-evolving proteins in higher plants. Photosynth Res 93: 89–100

Suorsa M, Rossi F, Tadini L, Labs M, Colombo M, Jahns P, Kater MMM, Leister D, Finazzi G, Aro EM, et al (2016) PGR5-PGRL1-Dependent Cyclic Electron Transport Modulates Linear Electron Transport Rate in Arabidopsis thaliana. Mol Plant 9: 271–288

Tadini L, Jeran N, Domingo G, Zambelli F, Masiero S, Calabritto A, Costantini E, Forlani S, Marsoni M, Briani F, et al (2023) Perturbation of protein homeostasis brings plastids at the crossroad between repair and dismantling. PLoS Genet 19: e1010344

Tadini L, Jeran N, Peracchio C, Masiero S, Colombo M & Pesaresi P (2020a) The plastid transcription machinery and its coordination with the expression of nuclear genome: Plastid-Encoded Polymerase, Nuclear-Encoded Polymerase and the Genomes Uncoupled 1-mediated retrograde communication. Philosophical Transactions of the Royal Society B: Biological Sciences 375: 20190399

Tadini L, Jeran N & Pesaresi P (2020b) GUN1 and Plastid RNA Metabolism: Learning from Genetics. Cells 9: 2307

Tadini L, Peracchio C, Trotta A, Colombo M, Mancini I, Jeran N, Costa A, Faoro F, Marsoni M, Vannini C, et al (2020c) GUN1 influences the accumulation of NEP-dependent transcripts and chloroplast protein import in Arabidopsis cotyledons upon perturbation of chloroplast protein homeostasis. The Plant Journal 101: 1198–1220

Tadini L, Pesaresi P, Kleine T, Rossi F, Guljamow A, Sommer F, Mühlhaus T, Schroda M, Masiero S, Pribil M, et al (2016) GUN1 Controls Accumulation of the Plastid Ribosomal Protein S1 at the Protein Level and Interacts with Proteins Involved in Plastid Protein Homeostasis. Plant Physiol 170: 1817–30

Tan X, Li K, Wang Z, Zhu K, Tan X & Cao J (2019) A Review of Plant Vacuoles: Formation, Located Proteins, and Functions. Plants 8: 327

Veciana N, Martín G, Leivar P & Monte E (2022) BBX16 mediates the repression of seedling photomorphogenesis downstream of the GUN1/GLK1 module during retrograde signalling. New Phytologist 234: 93–106

Vella D, Zoppis I, Mauri G, Mauri P & Di Silvestre D (2017) From protein-protein interactions to protein co-expression networks: a new perspective to evaluate large-scale proteomic data. EURASIP J Bioinform Syst Biol 2017: 6

Vera-Vives A, Perin G & Morosinotto T (2022) The robustness of NextGen-O2k for building PI curves in microalgae Padua

Vigani G, Di Silvestre D, Agresta AM, Donnini S, Mauri P, Gehl C, Bittner F & Murgia I (2017) Molybdenum and iron mutually impact their homeostasis in cucumber (*Cucumis sativus*) plants. New Phytologist 213: 1222–1241

Vogel MO, Moore M, König K, Pecher P, Alsharafa K, Lee J & Dietz K-J (2014) Fast Retrograde Signaling in Response to High Light Involves Metabolite Export, MITOGEN-ACTIVATED PROTEIN KINASE6, and AP2/ERF Transcription Factors in *Arabidopsis*. Plant Cell 26: 1151–1165

Wang J, Xia H, Zhao SZ, Hou L, Zhao CZ, Ma C Le, Wang XJ & Li PC (2018) A role of GUNs-Involved retrograde signaling in regulating Acetyl-CoA carboxylase 2 in Arabidopsis. Biochem Biophys Res Commun 505: 712–719

Wang S & Blumwald E (2014) Stress-Induced Chloroplast Degradation in *Arabidopsis* Is Regulated via a Process Independent of Autophagy and Senescence-Associated Vacuoles. Plant Cell 26: 4875–4888

Wang X & Auwerx J (2017) Systems Phytohormone Responses to Mitochondrial Proteotoxic Stress. Mol Cell 68: 540–551.e5

Wang Y, Selinski J, Mao C, Zhu Y, Berkowitz O & Whelan J (2020) Linking mitochondrial and chloroplast retrograde signalling in plants. Philosophical Transactions of the Royal Society B: Biological Sciences 375: 20190410

Wang Z, Gu C, Colby T, Shindo T, Balamurugan R, Waldmann H, Kaiser M & van der Hoorn RAL (2008) β-Lactone probes identify a papain-like peptide ligase in Arabidopsis thaliana. Nat Chem Biol 4: 557–563

Wiśniewski JR (2018) Filter-Aided Sample Preparation for Proteome Analysis. In pp 3–10.

Woodson JD (2022) Control of chloroplast degradation and cell death in response to stress. Trends Biochem Sci

Woodson JD, Joens MS, Sinson AB, Gilkerson J, Salomé PA, Weigel D, Fitzpatrick JA & Chory J (2015) Ubiquitin facilitates a quality-control pathway that removes damaged chloroplasts. Science (1979) 350: 450–454

Woodson JD, Perez-Ruiz JM & Chory J (2011) Heme synthesis by plastid ferrochelatase i regulates nuclear gene expression in plants. Current Biology 21: 897–903

Wu G-Z & Bock R (2021) GUN control in retrograde signaling: How GENOMES UNCOUPLED proteins adjust nuclear gene expression to plastid biogenesis. Plant Cell 33: 457–474

Wu GZ, Chalvin C, Hoelscher M, Meyer EH, Wu XN & Bock R (2018) Control of retrograde signaling by rapid turnover of GENOMES UNCOUPLED1. Plant Physiol 176: 2472–2495

Wu GZ, Meyer EH, Richter AS, Schuster M, Ling Q, Schöttler MA, Walther D, Zoschke R, Grimm B, Jarvis RP, et al (2019a) Control of retrograde signalling by protein import and cytosolic folding stress. Nat Plants 5: 525–538

Wu G-Z, Meyer EH, Wu S & Bock R (2019b) Extensive Posttranscriptional Regulation of Nuclear Gene Expression by Plastid Retrograde Signals. Plant Physiol 180: 2034–2048

Wulff A, Oliveira HC, Saviani EE & Salgado I (2009) Nitrite reduction and superoxide-dependent nitric oxide degradation by Arabidopsis mitochondria: Influence of external NAD(P)H dehydrogenases and alternative oxidase in the control of nitric oxide levels. Nitric Oxide 21: 132–139

Wurzinger B, Nukarinen E, Nägele T, Weckwerth W & Teige M (2018) The SnRK1 Kinase as Central Mediator of Energy Signaling between Different Organelles. Plant Physiol 176: 1085–1094

Xiao Y, Savchenko T, Baidoo EEK, Chehab WE, Hayden DM, Tolstikov V, Corwin JA, Kliebenstein DJ, Keasling JD & Dehesh K (2012) Retrograde Signaling by the Plastidial Metabolite MEcPP Regulates Expression of Nuclear Stress-Response Genes. Cell 149: 1525–1535

Xu L, Carrie C, Law SR, Murcha MW & Whelan J (2013) Acquisition, Conservation, and Loss of Dual-Targeted Proteins in Land Plants. Plant Physiol 161: 644–662

Yang Z, Lu S, Wang M, Bi D, Sun L, Zhou S, Song Z & Liu J (2014) A plasma membrane-tethered transcription factor, NAC 062/ ANAC 062/ NTL 6, mediates the unfolded protein response in Arabidopsis. The Plant Journal 79: 1033–1043

Yi X, McChargue M, Laborde S, Frankel LK & Bricker TM (2005) The Manganese-stabilizing Protein Is Required for Photosystem II Assembly/Stability and Photoautotrophy in Higher Plants. Journal of Biological Chemistry 280: 16170–16174

Yoo S-D, Cho Y-H & Sheen J (2007) Arabidopsis mesophyll protoplasts: a versatile cell system for transient gene expression analysis. Nat Protoc 2: 1565–1572

Zhao L, Peng T, Chen C-Y, Ji R, Gu D, Li T, Zhang D, Tu Y-T, Wu K & Liu X (2019a) HY5 Interacts with the Histone Deacetylase HDA15 to Repress Hypocotyl Cell Elongation in Photomorphogenesis. Plant Physiol 180: 1450–1466

Zhao Q & Meier I (2011) Identification and characterization of the *Arabidopsis* FG-repeat nucleoporin Nup62. Plant Signal Behav 6: 330–334

Zhao X, Huang J & Chory J (2018) Genome uncoupled1 mutants are hypersensitive to norflurazon and lincomycin. Plant Physiol 178: 960–964

Zhao X, Huang J & Chory J (2019b) GUN1 interacts with MORF2 to regulate plastid RNA editing during retrograde signaling. Proc Natl Acad Sci U S A 116: 10162–10167

Zhuang X & Jiang L (2019) Chloroplast degradation: Multiple routes into the vacuole. Front Plant Sci 10: 359

